# Environment-dependent evolution drives divergent adaptive strategies balancing parasite costs and competitive benefits

**DOI:** 10.64898/2025.12.17.694638

**Authors:** Teodora Simona Lungu, Louise Badruna, Olaya Rendueles

## Abstract

Prophages, phage genomes integrated into bacterial chromosomes, are widespread, yet how their costs to individual bacterial cells are balanced against benefits to the bacterial lineage remains poorly understood. This trade-off is particularly relevant in polymicrobial environments when prophage induction kills the host cell but releases phages that suppress competing bacteria. Here, to investigate how this conflict shapes bacterial and phage evolution, we experimentally evolved one polylysogenic strain in competition with a phage-sensitive strain across environmental conditions that modulate phage activity. By tracking bacterial growth and phage production over time, we uncover divergent, environment-dependent adaptive strategies consistent with a trade-off between direct costs of prophage activity and lineage-level (indirect) benefits of competitor suppression. At physiological levels of phage production, irrespective of the initial benefit, all populations evolved a 1,000-fold increase in phage production. However, under exacerbated phage production where induction imposed severe fitness costs, hosts purged prophages and adapted through accelerated bacterial growth. Phage production was negatively associated with bacterial yield, while competitive fitness showed a nonlinear relationship with phage production, with an optimum at intermediate-high phage activity. Whole-genome sequencing revealed that adaptation was accompanied by recurrent defects in cellular segmentation arising from mutations in the divisome and shifts in lysate composition, alongside the loss of plasmid-encoded antibiotic resistance genes. Together, our results reveal how resident genetic parasites can generate social trade-offs between the direct cost of prophage-induced death and the lineage-level competitive benefits. Our results provide a framework to study mobile genetic element dynamics within a kin selection mechanistic framework.

## INTRODUCTION

Competition is rampant in the microbial world^1,2^. Bacteria deploy a diverse array of competitive strategies, including chemical mechanisms (e.g., antibiotic and toxin production) ^3,4^ and biological mechanisms such as contact-dependent killing mediated by Type VI secretion systems or other CDI systems^4^, as well as metabolic forms of interference like metal scavenging or pH modulation^5^. Another strategy to attack conspecifics, which are expected to have overlapping ecological niches and thus be in direct competition for the same resources ^6–8^, relies on the release of infectious phages. Bacteriophages are very abundant predators of bacteria. Nearly half of the sequenced bacterial genomes are lysogens^9^, that is, they have phages inserted in their genome. Although such bacteriophages (hereafter referred to as prophages) can provide context-dependent benefits, they are evolutionarily parasitic elements whose persistence reflects a balance between costs to the host and benefits to phage transmission. Upon induction, prophages excise from the host’s genome, replicate within bacterial cells and kill its host to release infectious virions. Consequently, prophage activation imposes the ultimate cost on individual bacterial cells (death), while the released virions can kill susceptible competitors, providing an indirect advantage to clonally-related lysogens (kin).

Kin selection theory provides a framework for how individually costly strategies can be maintained when indirect (lineage-level) benefits to clonally related cells outweigh direct costs^10,11^. However, because prophages are both mobile genetic elements and potential social weapons^12^, prophage maintenance reflects an evolutionary interplay between parasite transmission strategies and host inclusive fitness. While theoretical models demonstrate that investment in costly, or lethal, strategies, like toxin release, can be favored under facultative stress or competitor pressure^13^, how lysogenic bacteria adapt across environments that differ in the cost–benefit ratio of phage production remains poorly understood. We hypothesize that evolutionary outcome should therefore depend on the balance between the direct cost of prophage induction and its indirect benefit through competitor suppression.

To investigate how ecological conditions shape social trade-offs, we set-up a coevolution experiment using in a polylysogenic, multi-drug resistant *K. pneumoniae* (ST14) cocultured with a phage-sensitive competitor (hypervirulent *K. pneumoniae* BJ1). Our previous work analyzed the different phage resistance mechanisms of the phage-sensitive BJ1 across different environments. We showed that the fitness costs of the different phage-resistant strategies drove frequency of resistant clones through time. Contrary to the expectation, we showed that lysogenization (integrating the phage in the genome) which had been a resistance mechanism observed when the strain BJ1 was attacked by purified phage from ST14^14^ was not an efficient strategy in the presence of ST14 itself, due to the cost of spontaneous phage excision from newly lysogenized BJ1^15^. This highlighted how very small changes in ecological complexity, which increases lineage costs, can alter evolutionary outcomes within microbial communities due to the presence of multiple selective pressures (i.e. cost of phage resistance and presence of a competitor).

Here, we took advantage of this coevolution experiment to ask whether selection favors investment in phage production providing a lineage-level benefit, despite the individual costs, and whether there is a point in which phage activity becomes excessively costly. The mixed population coevolved under three different ecological conditions resulting in different degrees of phage infection, and thereby different points along the cost–benefit gradient^15^; (i) LB representing some individual cost and some degree of lineage-level benefit (see Methods), (ii) LB supplemented with 0.2% citrate to inhibit phage infection due to calcium chelation^16^, whilst still paying the individual cost and (iii) LB with mitomycin C (MMC, 0.1 μg/mL) to favor phage induction^15,17^, and maximizing the benefits, but also the cost. We expected evolutionary outcomes to depend non-linearly on the cost-benefit ratio of phage activity. Specifically, we predicted that when phage infectivity is blocked (citrate), selection should favor reduced phage activity and increased bacterial growth. Under excessive phage induction (MMC), the severe cost of host cell death should favorise individual-level strategies, thereby reducing phage induction rate^18^, accumulating deleterious mutations in the phage^19^ or favor prophage loss or inactivation^20,21^. At intermediate levels (LB), phage production should yield an optimal competitive benefit that offsets host death. Our findings revealed the emergence of two divergent strategies driven by environmental conditions. Whole genome sequencing and experimental validation allowed us to show that populations adapted by defects in capsulation and cell division, as well as strong reduction of antibiotic resistance. Analyses of phage sequences and frequencies in end-point lysates show shifts in lysate composition with potential implications in evolutionary outcomes. Taken together, sor work highlights how social trade-offs and pleiotropic by-products shape bacterial adaptation, prophage stability and maintenance of antibiotic resistance genes in multidrug-resistant pathogens.

## RESULTS

### Environment conditions select for divergent bacterial growth strategies

In a previous experiment, we allowed three independent cocultures of *K. pneumoniae* polylysogenic strain ST14 to coevolve with the phage sensitive strain *K. pneumoniae* BJ1^15^ in three environmental conditions. Each population was transferred daily (1%) during 30 days, accounting for ca 200 generations. Three clones of the lysogen ST14 were isolated from each evolving population throughout the experiment (day 10, 20 and 30). This results in a total of 81 independent clones. To evaluate how selection acts on individual-level fitness, and whether evolved clones adapted by prioritizing direct cell reproduction (growth yield), we grew independent clones in their respective evolutionary environment across the three environmental regimes. We then calculated maximum population yield and the area under the growth curve which takes into account growth rate, lag time and yield. Overall, we observe that clones isolated from populations evolving in MMC increase growth, as shown by the area under the curve (AUC). This enhanced growth is mostly driven by a higher population yield (Figure 1). To statistically compare changes across evolutionary treatments whilst circumventing inherent differences due to the environment, all measures were also calculated as relative to the ancestor’s performance in each environment (µ=1) (Figure S1, Table S1). Unexpectedly, clones from populations having evolved in citrate, without phage infection, and in LB, do not outperform their ancestor, and some tend to grow worse (Figure 1 and Figure S1). Statistics corroborate that changes in yield (and AUC) were dependent on environmental phage pressure, namely, populations evolving in MMC grow better, whereas all others did not change or performed worse when compared to the ancestor (Figure S1). These same trends were independent of the day in which clones were isolated, revealing that these adaptations happened within the first ten days (Figure S1). We also tested for differences in generation time and, on average, on the 27 end-point clones no significant changes were observed (µ=1.02, Wilcoxon test, *P*=0.34, difference from 1), irrespective of the evolutionary environment (Kruskal-Wallis, *χ*^2^ = 1.4, *P*=0.4, Figure S2A).

**Figure 1.**
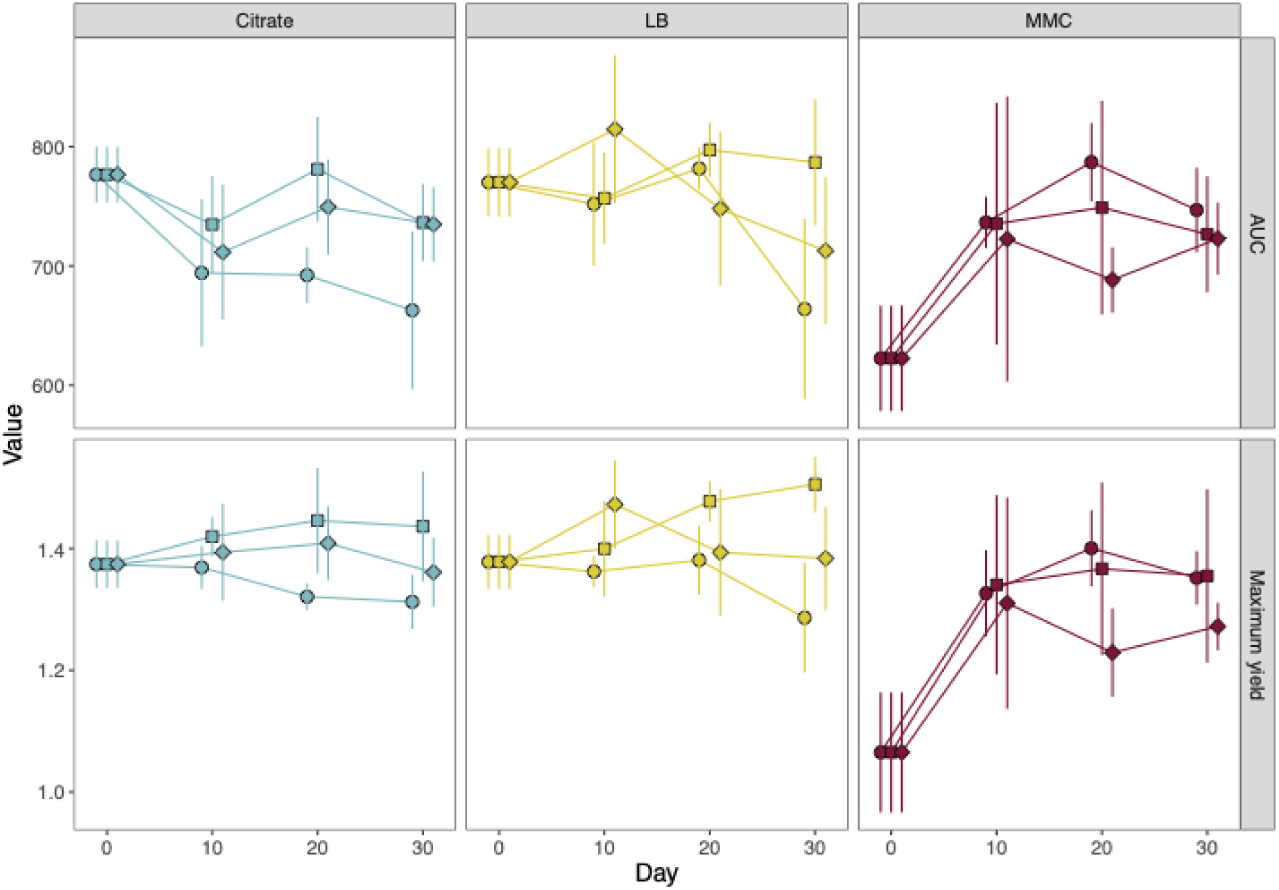
Growth of independent clones isolated from each population at different time points. Growth is assessed both by the area under the curve (AUC, upper panels) and the maximum yield of each individual clone of each population (lower panels). Growth was measured in the environmental conditions in which each population evolved. Each point represents the average of three independent clones (N=81), which were tested in at least three biological replicates. Different shapes represent different populations. Error bars correspond to the standard deviation of the mean. More details and statistics are provided in Figure S1 and Table S1.

Finally, we tested whether the observed increase in yield in MMC-evolved populations would fully compensate the cost of cell death induced by the phage burst. To test this, we compared the maximum yield of clones evolved in MMC to that of the ancestral strain grown without MMC, i.e. in LB. Overall, statistical analyses revealed very marginal differences between the evolved clones and the ancestor. This suggests that the evolved clones in MMC offset the cost of phage induction as they reached similar levels than the ancestor in the absence of phage induction (Figure S2B). A similar pattern was observed when considering the area under the curve (AUC) (Figure S2C). Interestingly, among the 27 clones isolated from the MMC environment, eight (∼30%) show higher yields than the ancestor, with five of these increases being statistically significant (Figure S2B, dots above the dashed line), suggesting that growth increases of these evolved clones totally compensate and improve ancestral performance despite the use of mitomycin C.

Taken together, our results show that changes in growth are contingent on evolutionary treatment and are linked to phage infectivity, that is, in environments in which phage burst results in exacerbated population death (MMC), yield of evolved clones increase. On the contrary, clones from populations in which phage infection is blocked or phages are released at physiological levels, they do not improve growth, with many exhibiting reduced growth rates, suggestive of trade-offs between individual cell growth and lineage-level benefits.

### Phage production imposes direct individual costs but yields an intermediate competitive optimum

In our evolutionary experiment, adaptation could be achieved either by directly maximizing individual growth or by investing in phage production which would indirectly benefit kin. To test this, we assessed phage production of each population at different time points. We selected one clone from each population and each time point and grew them in their own evolutionary conditions. We then filter-sterilized all lysates and added them to ancestral prey (1:100) and measured growth. Our results show that prey population grew much worse in the presence of lysates from evolved clones, with an exception, those grown in the presence of lysate from MMC-evolved populations (Figure 2A). This suggested that clones evolved in MMC produced less phages or that these could not infect. The opposite would be true for clones evolved in other environmental conditions, that is, either evolved clones were able to produce more phage or they evolved a new mechanism to limit growth of ancestral prey.

**Figure 2.**
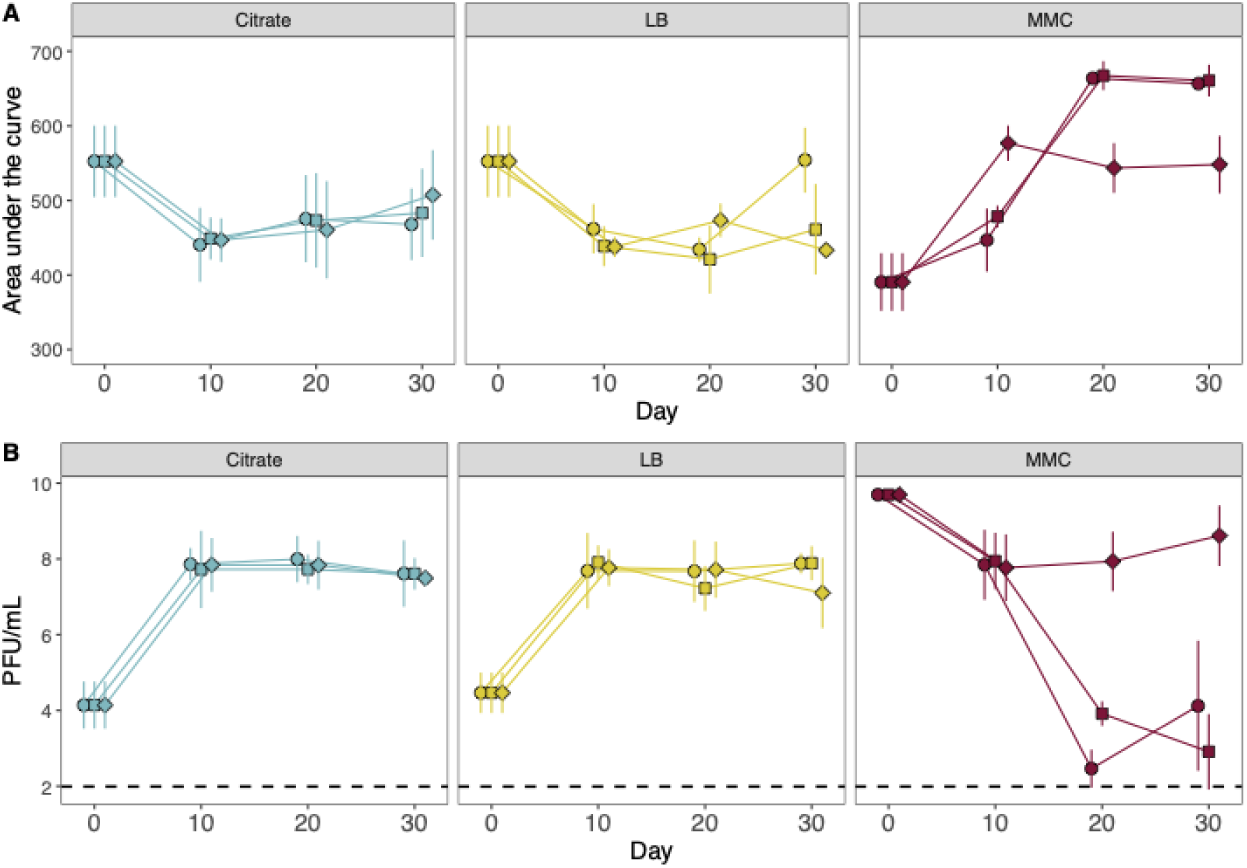
Evolution rapidly shifts phage production in opposite directions. **A**. Phage was evaluated by their ability to attack their prey (strain BJ1) in their evolutionary conditions (liquid shaking). Results are presented as the area under the growth curve of BJ1 in the presence of unpurified lysates produced by a representative clone of each population at each time point. Decrease in the AUC represents increased death of BJ1. Different shapes represent different populations. Error bars correspond to the standard deviation of the mean. At least three independent biological replicates are shown. **B.** Phage quantification of the different unpurified lysates produced by a representative clone of each population on a lawn of BJ1 is represented as the log_10_ of the plaque forming units per mL (PFU/mL). The dashed line represents the limit of detection of our assay. Relative changes and statistics are presented in Figure S3.

To distinguish these two possibilities, we quantified phages in each lysate by counting the plaques on a lawn of ancestral prey. In agreement with bacteria-lysate cocultures, we observed that clones evolving in citrate and in LB very rapidly increased phage production. On average, they exhibited ca. 3 logs more phage production in the first ten days, and this remained stable for the rest of the experiment (Figure 2B and Figure S3). Clones evolving in MMC at first showed ca 2-logs reduced phage infection but this declined more at later stages, near to the limit of detection of our assay in two of the three populations.

Our results showed a clear trade-off between phage production (Figure 2B) and yield (Figure 1A). Indeed, relative changes in phage production showed a strong negative correlation with relative bacterial yield (Figure 3A; GLM,P = 0.0003). This trade-off split the evolved populations into two distinct adaptive strategies: clones either invested in exacerbated phage production despite reduced growth, or maximized individual growth while minimizing phage release. Further, this trade-off was established very early during the evolution experiment. Closer analyses of the intermediate time points suggest that populations adapted in a step-wise manner with large changes within the first ten days in phage production, followed by a second step in which growth was optimized (Figure S4).

**Figure 3.**
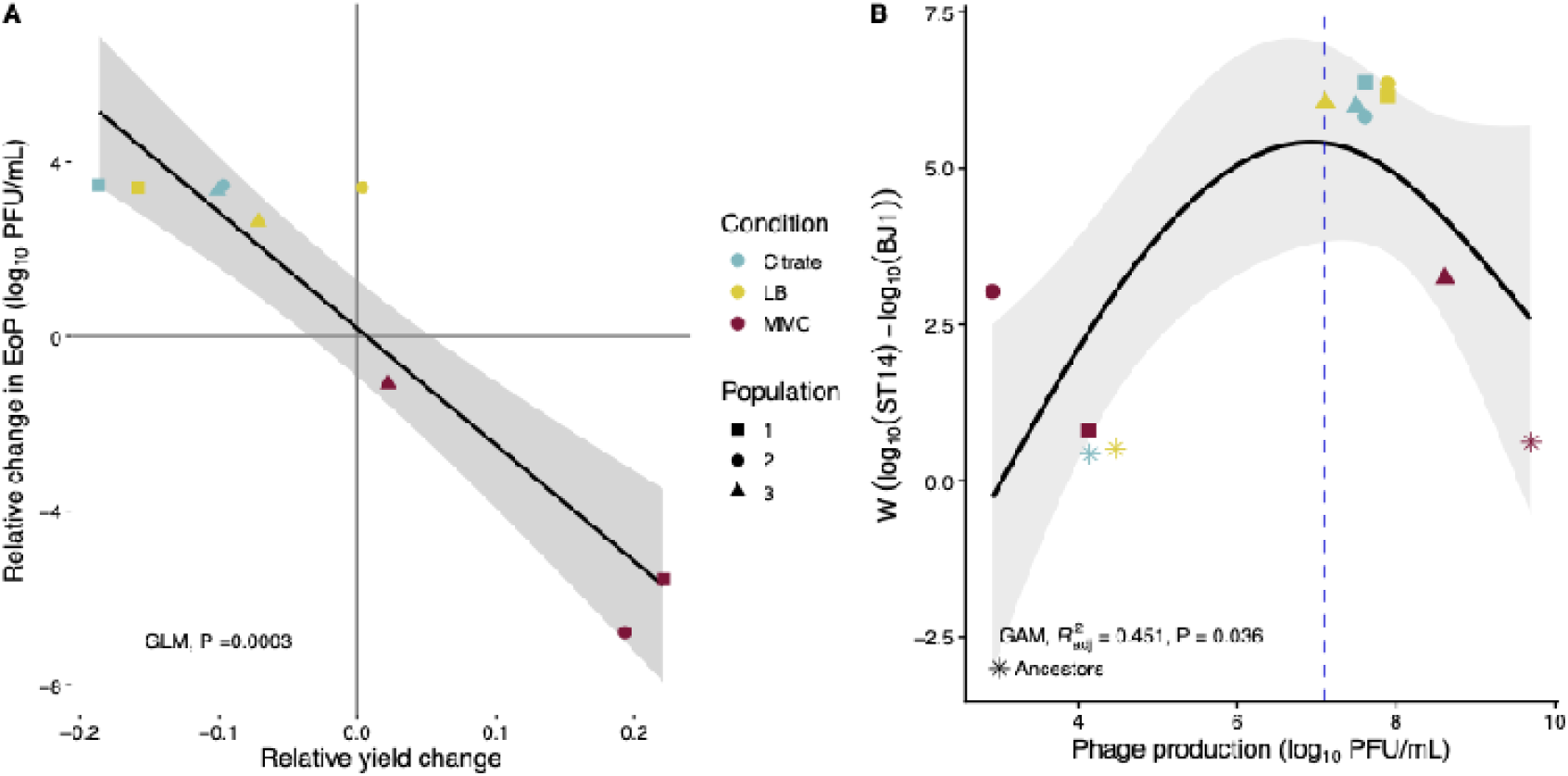
Phage production is costly but generates a nonlinear competitive benefit. **A.** Negative correlation between the change in phage production and growth, relative to their common ancestor. Phage production was measured by PFU/mL, and represented in a log_10_-transformed scale. Inclusion of all intermediary data (day 10 and 20) does not qualitatively alter the correlation. The black line represents a generalized linear model (GLM) and grey shading corresponds to the standard error. Colors indicate the environmental condition and point shapes indicate population. **B**. Relationship between endpoint phage production and competitive fitness, as calculated by the proportion of each lineage in the endpoint populations^15^. Points show evolved clones, with symbols corresponding to environmental condition. Stars represent the ancestral population. The solid line shows the fitted generalized additive model (GAM) and the corresponding confidence interval. The curve indicates an apparent optimum in competitive fitness at intermediate-high phage production.

We next asked whether the direct cost of phage production was offset by a lineage-level competitive benefit (*W*) (Figure 3B). Fitness showed a non-linear, hump-shaped relationship with phage production (GAM, 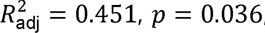, variance explained = 54.4%), indicating an intermediate peak in competitive fitness rather than a strictly monotonic benefit. While bootstrapping demonstrates variance around the precise optimum (6.9 log10 PFU/mL), as expected from the limited number of evolved populations, the non-linear trajectory confirms intermediate-to high phage production maximizes fitness before host death costs predominate. Evaluating ST14 density alone (without competition) yielded a lower optimum (6.3 log_10_ PFU/mL), suggesting that competitor killing pulls the evolutionary optimum toward higher phage release.

Altogether, these patterns are consistent with an evolutionary trade-off in which phage production is favored when its lineage-level benefit through competitor suppression offsets its direct cost to the producing cell, thereby fitting a kin selection framework.

### Evolved phage–prey interactions reveal density-mediated infection and expanded host range

One of our original expectations was that populations evolving in citrate would adapt by maximizing individual growth, as lineage-level benefits were, a priori, not possible, due to calcium chelation blocking infection. Yet, this was not the case. In our previous study, we had observed that some phages evolving in citrate could lysogenize the prey BJ1^14,15^, suggesting some phages might overcome citrate-mediated protection. However, purified lysates from endpoint citrate-evolved ST14 remained unable to infect prey at ancestral production levels (MOI of 0.01, Figure S5A). In contrast, infection occurred when crude evolved lysates, and thus much higher phage concentrations were used (Figure S5B). Consistent with this, increasing phage concentration overcame citrate-mediated protection, with prey growth inhibited at high (evolved) but not low (ancestral) MOIs (Figure 4A). Thus, citrate did not prevent the lineage-level benefit, once phage excision levels increased to a sufficient high density and infect previously protected prey.

**Figure 4.**
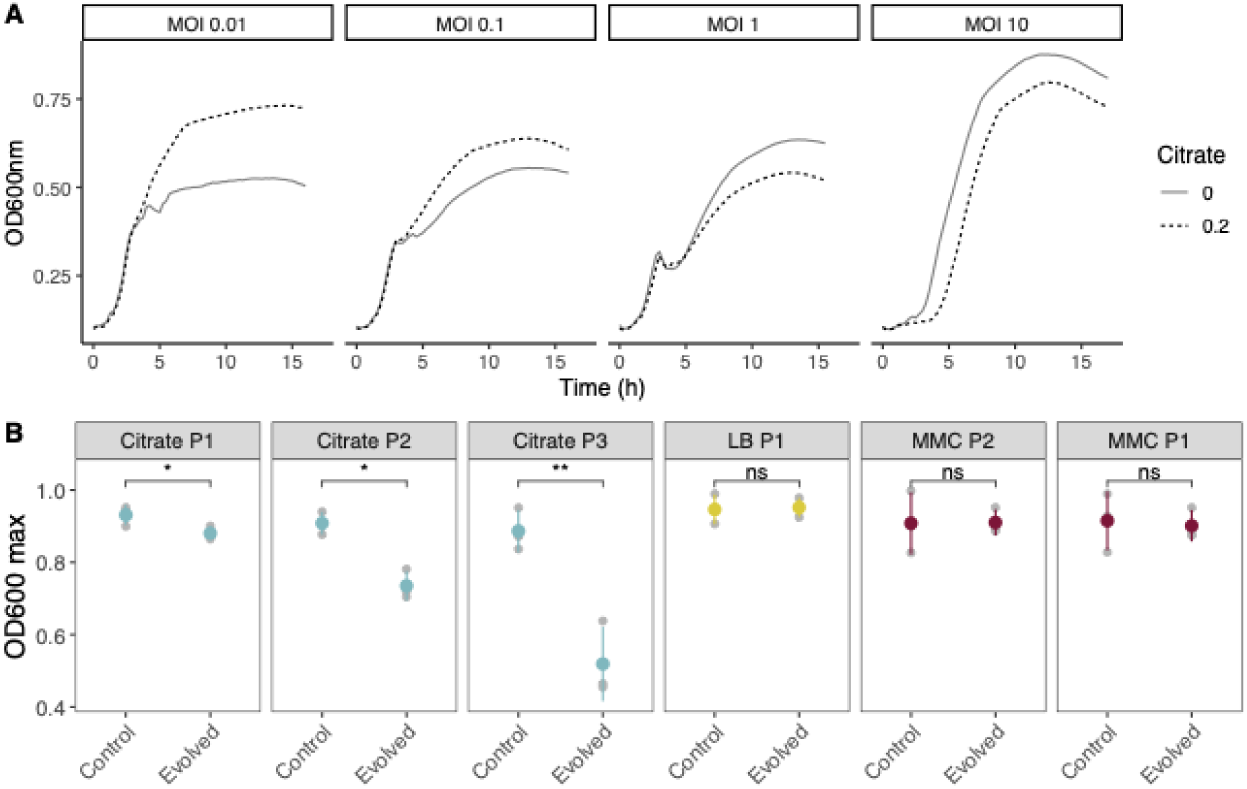
Evolution of phage infection patterns. **A.** Growth curves of prey *Kpn* BJ1 when ancestral PEG-precipitated lysates were added at different initial multiplicity of infections (MOI). Infection was tested in media with (dashed lines) or without 0.2% citrate (solid lines). Error bars are not included for visibility purposes. Experiments were performed in triplicate, and the average is shown. **B.** The ability of end-point lysates to infect resistant clones, which emerged during the first days of the experiments, was assessed in evolutionary conditions. Unpurified lysates were tested alongside the control condition (plain media). Each resistant clone was challenged against a lysate from a representative evolved ST14 clone from the same population (indicated on top). The maximum absorbance (OD600nm max) reached by each culture is shown. Data for the area under the curve is shown in supplementary Figure S5C. Statistics correspond to pairwise t-tests (ns p-value > 0.05, * p-value <0.05, ** p-value <0.01).

We also hypothesized that phages and their prey would engage in a coevolutionary arms. Consistent with this, we expected that end-point lysates could infect phage-resistant clones isolated early in the experiment (within the first nine days). To test this, we added crude lysates to growing cultures of selected several lightly capsulated resistant clones previously described^15^ from the same population as the phage-producer. Two of the resistant clones evolved in citrate (P1 and P2) have mutations in genes necessary for capsule production (*wcaJ*) and tethering to the outer membrane (*wzi*)^15^. Clones isolated from LB and MMC all had mutations in the *orf13* gene of the capsule, an acetyltransferase expected to alter the chemical composition but not the amount of the capsule. All resistant clones but one evolved in citrate (P3) inhibit adsorption^15^. Our results show that the three end-point lysates from ST14 evolved in citrate tested could infect BJ1 resistant clones (Figure 4B and Figure S5C), whereas those with mutations in *orf13*, they remained resistant. Control experiments with non-capsulated clones reveal that these were all resistant to evolved lysate (i.e. no PFUs were observed on a lawn of non-capsulated prey).

Altogether, our results indicate that while phages evolved in citrate did not *sensu stricto* bypass citrate blocking, evolved interactions allow them to infect previously resistant hosts suggesting an arms race dynamic.

### Striking parallel chromosomal mutations reveal common changes in capsule production and cell segmentation

To underpin the molecular mechanisms of adaptation of the polylysogen, we performed paired-end short-read whole-genome sequencing of one end-point clone (day 30) from each population and compared the sequence against that of the ST14 ancestor. All chromosomal mutations in bacterial genes are presented in Table S2. Despite the divergent evolutionary strategies observed across environments, sequencing revealed some remarkable examples of parallel convergence (Figure 5A). All sequenced clones had at least one mutation in *wcaJ*, the first gene in the biosynthesis cascade of exopolysaccharidic capsule, leading to arrest of capsule production. Uronic acid quantifications of capsule production confirmed unencapsulation as soon as day 10, matching our previous visual observations^15^ (Figure 5B).

**Figure 5.**
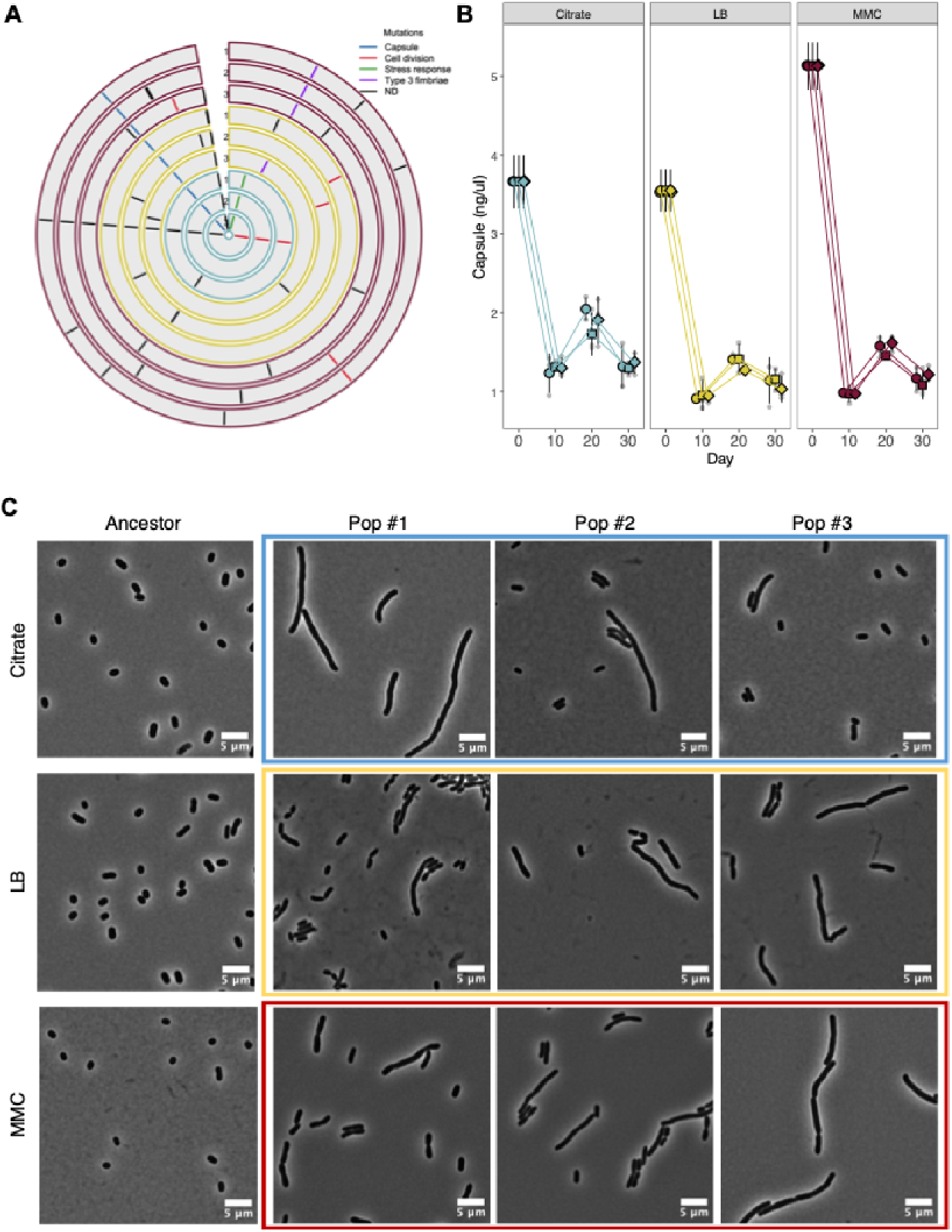
Mutations identified in a representative clone of each end-point population lead to capsule reduction and defective cell segmentation. **A.** Mutations were detected by *breseq* and *snippy* and detailed in Table S2. Each line represents a mutation, colored according to the pathways most frequently targeted across evolved clones. ND bins together all other mutations. Visualization was done with the R package *circlize.* B. Capsule production of each individual end-point clone in its own evolutionary condition, as determined by the uronic acid method. Each grey point represents the average of three independent replicates for each of the three clones tested from the population (N=81 clones). The slight increase observed at day 20 is a batch effect caused by differences in experimental timing and the use of a fresh batch of reagents, which produced a brighter signal. This is also reflected in the error bars of ancestral capsule production. **C.** Representative micrographs on agar pads of sequenced ST14 end-point clones were taken. Overnight cultures were grown in LB and 0.6 µL of culture were deposited on a 1% agar pad on a microscope slide. Images were taken on an inverted microscope at a magnification of x100. Scale bar, 5 µm.

Similarly, each sequenced clone, presented mutations in central players of the cell divisome, mainly *ftsX* and *ftsW*, *zipA* and *envC*. Mutations in these genes generally lead to defective hydrolysis of the septum^22,23^. Interestingly, these mutations were not observed in populations evolving in citrate. However, in these populations, we observe mutations in *tatC*. The Tat translocation system is in charge of the translocation of folded proteins across the inner membrane, and notably of AmiA and AmiC, two amidases which are critical for cell separation in *E. coli* ^24^. To confirm this, we examined the sequenced clones from each population using phase-contrast microscopy. All clones, independently of their evolutionary treatment, showed dramatic deficiencies in cell division and striking cell elongation (Figure 5C). These mutations appear early during the evolution experiment, as clones isolated at day 10, already presented segmentation defects (Figure S6). Finally, we also observed deletions in all populations evolving in mitomycin C and one in LB of the intergenic region just upstream *mrkH*, a major regulator of fimbriae^25^. We hypothesize that fimbriae production is most likely counterselected due to the culture agitation.

Taken together, and despite the marked differences in adaptive strategies, we observe striking genetic parallelism leading to reduced capsule production and cell segmentation defects, suggesting common bacterial responses to phage-mediated selection.

### The cost of phage activity reshapes the resident prophage community

The polylysogenic strain ST14 carries 8 different phages or phage remnants, five of which are chromosomal and the other three are located in its plasmids (Figure 6A and Table S3). To understand the different patterns of phage infection in the evolved populations, we analyzed all mutations and genome reorganizations which occurred in regions identified as phages (Figure 6A). Similar to the changes in the bacterial genes, we saw a high degree of parallel evolution. In all end-point clones two insertions in a homopolymeric track of 6 purines coding for tail fiber proteins in phages #3 and #4 (Table S4). Phage #4 also carried a ∼5-kb inversion downstream of a DNA invertase, comprising five genes, including three involved in fiber protein synthesis (Figure S7). However, sequencing coverage indicated only minimal changes in the copy number of phages #3 and #4 (Figure S8A).

**Figure 6.**
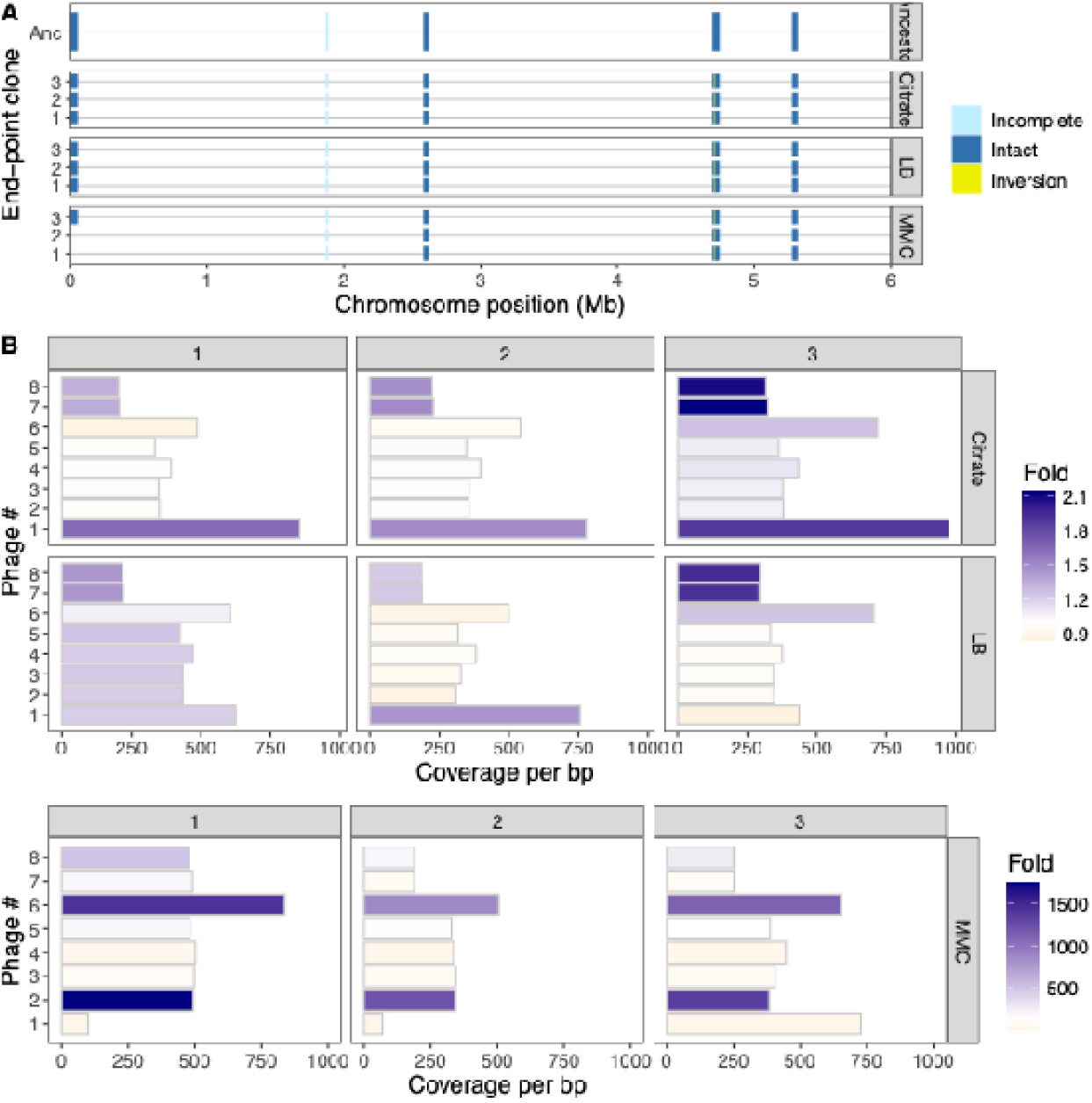
Phage evolution across end-point evolved clones. **A.** Location and size of the five phages in the chromosome, four of which are considered intact and one incomplete. The image illustrates the ancestral sequences as well as the end-point evolved clones. Details of mutations in phage regions are presented in Table S4. **B.** Illumina read coverage per base pair across phage genomes for each end-point clone. Fold change compared to the ancestor was calculated by dividing the total raw reads of each phage in evolved clones and ancestral lysates. Raw ancestral reads in the presence and absence of MMC are shown in Figure S8.

The most striking changes occurred in MMC-evolved populations. Two endpoint clones carried a large deletion spanning phage #1, which was the predominant phage in the ancestral MMC lysate. These clones exhibited nearly undetectable phage production, whereas clones from population #3 retained phage #1 and could still infect BJ1 at near-ancestral levels (Figures 2B and 6B). In this population, reduced induction of phage #1 could be associated with a nonsynonymous mutation in *hflX*, a gene implicated in regulating the stability of the phage cII protein and the lysogeny–lysis decision in lambda-like phages^26,27^. These changes suggest that when phage activity was highly costly, populations either lost phage #1 or reduced its induction.

Finally, mapping of raw reads revealed environment-dependent changes in the composition of evolved lysates. In MMC, the loss or reduction of phage #1 was accompanied by increased excision of phages #2 and #6. In LB and citrate, where phage production increased, phage #1 showed increased excision, with a consistent but minor contribution of plasmidic phages #7 and #8 (Figure S8B). In one LB population, all resident phages increased excision to similar extents. The different changes in phage abundance and excision suggest that interactions among co-resident phages may contribute to the final lysate composition.,

Overall, these analyses show that the environmental cost of phage activity also shaped the resident prophage community. When phage production was highly costly, the host lost or reduced induction of the dominant prophage, whereas under less costly conditions, selection favored increased phage production and shifts in lysate composition..

### Deletion of phage region result in reduced antibiotic resistance

The strain polylysogen ST14 used in this study is a multi-drug resistant *Klebsiella pneumoniae.* Phages #6 and #7 are located in the largest plasmid and encode for 13 different antibiotic resistance genes, from various classes, including fluoroquinolone and macrolide resistance genes (Figure 7A). Our sequencing data also revealed a strongly selected deletion within phage regions located in the plasmid (Table S4). The deleted regions overlapped with the predicted phages as well as with multiple genes involved in heavy metal resistance, i.e. mercury, and antibiotic-resistance. To experimentally validate whether the loss of these genes resulted in changes in antibiotic resistance, for each of the sequenced clones, we quantified the minimal inhibitory concentration (MIC) of two different antibiotics belonging to two different classes, namely erythromycin, a macrolide, and streptomycin, an aminoglycoside (Figure 7B). All evolved clones showed markedly reduced MIC values, with decreases ranging from two-fold to eight-fold.

**Figure 7.**
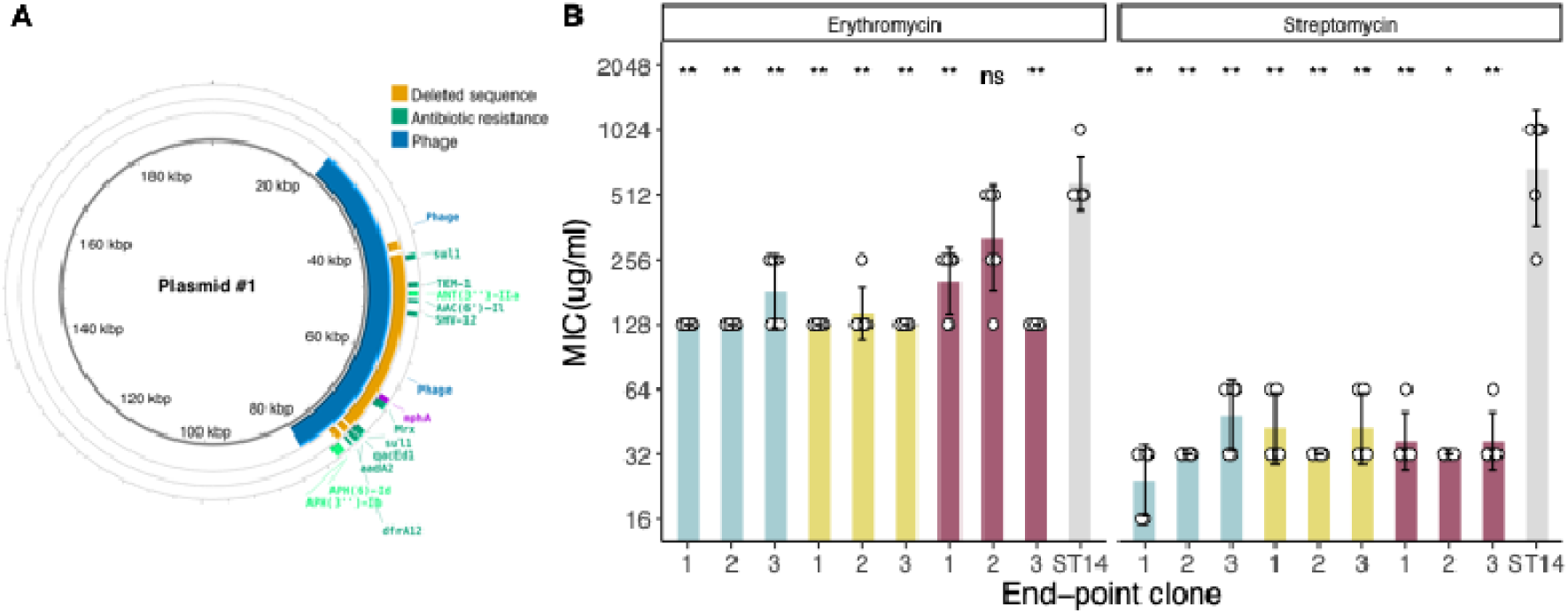
Loss of antibiotic resistance due to deletion in phage region. **A.** Map of plasmid was created using Proksee^28^. In blue, an two overlapping phage sequences were detected using Phastest 3.0^29^ (phage #6 and #7, Table S4). In yellow, areas that have been deleted in different end-point individual clones. The outer circle represents antibiotic resistance genes as identified by CARD v1.3.1^30^. *mphA* gene is highlighted in purple and typically associated to resistance to erythromycin. Genes highlighted in light green are associated to resistance to streptomycin. **B**. Minimal inhibitory concentration (MIC) of each sequenced end-point clone from each population. Each individual point represents an independent experiment. Points indicating an MIC of 1024 μg/mL, represent an MIC > 512. Bars in blue, yellow and red represent clones from populations evolved in citrate, LB and MMC, respectively. Wilcoxon tests for comparison against the ST14 ancestor (grey) were performed: ns p> 0.05, *p<0.05,**p<0.01.

Overall, our results reveal an adaptive trade-off between the competitive advantage provided by the phage and the genetic burden imposed by the cost of antibiotic resistance.

## DISCUSSION

Prophage induction creates a social conflict in which the individual bacterial cell pays the cost of phage production and lysis, while the resulting phage particles can suppress susceptible competitors and benefit surviving members of the same bacterial lineage (kin)^31–33^. Here, we used a coevolution experiment to examine how this trade-off shapes bacterial and resident phage evolution under distinct environmental conditions^15^. We found two divergent adaptive strategies: lysogens either invested in lineage-level benefits and increased phage production despite reduced bacterial yield, or increased individual growth and reduced phage production. Population dynamics at the end of the coevolution experiment showed a nonlinear relationship between competitive outcome and phage production, with an apparent optimum at intermediate-high phage activity. Our results are consistent with an evolutionary trade-off in which the lineage-level benefit of competitor suppression can offset the direct cost of phage production, fitting a kin-selection framework.

We had hypothesized that in environments with exacerbated phage production and lysis, strategies to control or downregulate the excessive burst could evolve^18,19,21^, in order to compensate for cell death, as has been previously observed in colicin-producing *E. coli*^34^. Clones from three populations rapidly reduced phage production, and in two of these populations the reduction reached levels close to the limit of detection of our assay (approximately a ≥7-log decrease relative to the ancestor). Whole genome sequencing showed the purge of one of the phages in two of the populations. The third population (population #3) had an intermediate phenotype, showing a modest reduction in phage production while also displaying a slight improvement in growth relative to the ancestor (Figure 3). Reduced phage production could be explained by the fixation of a mutation in *hflX*, which codes for a protein controlling the balance between lysogeny and lytic cycle^35^, which domesticated the phage most induced by MMC. Interestingly, phage production of this clone is similar to that of endpoint clones evolved in other environments, *ca* 10^8^ PFU/mL (Figure 2B and 3B). This suggests that there may be an upper optimal threshold of phage production beyond which growth costs become substantial. In our previous study, individual-based modelling suggested that phage production involving up to ∼5%, but less than 10%, of the population could be sustainable. Beyond this level, the competitive cost became too high^15^. Statistical modelling placed a competitive fitness peak at ca 6.9 log_10_ PFU/mL, whereas ST14 performance in the absence of competition showed a lower optimum of approximately 6.3 log_10_ PFU/mL. Due to the limited number of evolved populations analyzed, and substantial uncertainty in the precise location of these optimums derived from bootstrapping, we interpret the peak as evidence for an intermediate competitive optimum rather than as a precisely estimated phage-production target.

An important question is to what degree BJ1, the originally sensitive strain, is also directly or indirectly determining adaptation and phage titers? In the two conditions in which the polylysogen was allowed to produce phage at physiological levels very rapidly during the first ten days, phage production increased by several logs. This aligns with a strong reduction of phage-sensitive bacterium and an emergence of resistance by loss of phage receptor^15^. The order in which these events unfolded cannot be clearly established, but our data show that phage resistance emerged already during the first three days. We suspect that clones producing larger amounts of phage were selected afterwards to provoke lysis from without in the phage sensitive bacteria. Indeed, population dynamics of the originally sensitive bacteria showed a second decrease in population yield after ten days. This, alongside the fact that evolved phages can infect BJ1 clones (Figure 4B) resistant to ancestral lysate, is indicative of a coevolutionary arms race. However, in conditions in which the polylysogen was forced to produced non-physiological levels of phage, the sensitive strain dropped in frequency faster, but also resistance emerged faster^15^. More importantly, we did not observe a second drop in population yield of strain BJ1, most likely because there is little phage in the lysates, and thus less lysis from without or a more relaxed evolutionary arms race. These data collectively suggest that both the environment and evolution from the phage sensitive could determine the evolution of the polylysogen. However, this interpretation remains speculative, as higher temporal resolution in lysate activity and additional data at key time points would be required.

Despite the two different divergent strategies, all populations exhibited common phenotypic traits which may reduce the cost of phage exposure. Indeed, we had also shown that ST14 was somewhat sensitive to its own lysate^14^, even more so, when exposed to high titers of phage as those attained during the evolution experiment suggesting that mechanisms against superinfection are not optimal, or there is some lysis from without^36^. All 81 examined clones were non-capsulated, including the 27 clones isolated at day 10, due to mutations in the capsule operon. This outcome was expected as capsule loss in nutrient-rich media such as LB has been shown to reduce growth cost^37^. Yet,, since the capsule serves as the main receptor for lysogenic phages^14,38^,capsule inactivation may have also been selected as a by-product because it may protect ST14 from lysis from without. In a similar way, all end-point clones and some intermediary clones presented defects in cell segmentation. These may represent either a general adaptation to prolonged growth or an indirect consequence of selection under phage pressure. On the one side, cell segmentation defaults also emerged in another *K. pneumoniae* short-term evolution experiment in LB^39^ and in *P. fluorescens*^40^ in the absence of phages pressure. On the other hand, a recent study showed that mutations in *tatC* in non-encapsulated *Klebsiella,* confer infection resistance to a broad range of phages belonging to different families, though the precise mechanism remains to be determined^41^. We speculate that failure to complete segmentation during cell division could provide an extra advantage by reducing accessible surface area to phages, thereby also reducing the risk of superinfection. Overall, these parallel mutations may reflect selection not only by environmental conditions, but also by their capacity to mitigate phage pressure, for instance by limiting lysis from without and superinfection.

Beyond changes in bacterial genes, phages and lysate composition also evolved during the experiment. All isolated clones share a five-ORF inversion in phage #4. This inversion does not disrupt any ORF, and is not predicted to impact phage excision and production. Consistent with this interpretation, we observed only minor changes in the induction of phage #4 across populations. It may, however, attenuate production of fiber proteins, which could in turn impact host recognition^42^. Such a change may have been favored following changes in the bacterial capsule, the primary receptor for several phages in this system^14,38^. In contrast, phage #1 showed pronounced and environment-specific dynamics. Its purge in two MMC-evolved populations coincided with strong reduction of infection efficiency against the ancestral prey strain, suggesting that phage #1 is a major contributor to infectivity in ancestral MMC lysates. Accordingly, sequences of ancestral lysates revealed dominance of phage #1 and only trace levels of phage #4. However, phage loss may also have indirect effects, as recent studies indicate that phages engage in complex social interactions and can cooperate, compete and exploit other co-occurring phages^43^. The absence of phage #1 may therefore alter the induction, replication, or packaging efficiency of other prophages, reshaping overall lysate composition. In agreement with this possibility, we observed increased induction of phage #6 and of the incomplete phage #2 relative to ancestral levels, although this could not compensate the depletion of phage #1 in the lysate. We believe that these changes are unlikely to reflect a general loss of infectivity toward the ancestral host, as phage #2 has previously been shown to lysogenize BJ1^14,15^. This may reflect increased competition between phages. Indeed, while some of the changes in lysates followed a predictable additive pattern, epistatic interactions among phages could shape the final lysate composition by modifying the ecological and competitive landscape. Subtle but consistent differences in phage profiles among evolved lysates, even within the same environment (Figure 6B), support the idea that phage–phage interactions evolve during adaptation on mixed communities.

Finally, the evolutionary changes in phage content extended beyond phage activity itself. We observed recurrent loss of antibiotic-resistance genes located within prophage-associated plasmid regions, accompanied by reduced resistance to erythromycin and streptomycin. In the absence of positive selection, this loss is consistent with the cost of maintaining resistance ^44,45^. This loss mirrors a recent study where, as a byproduct of general adaptation, the loss of ARGs in a plasmid was observed and resulted in an increase in its capacity to across the population (both by vertical and by horizontal transfer)^46^. In *Kpn*, several studies have shown loss of ARGs. For example, the introduction of phage in a population resulted in a large plasmid deletion containing several resistance genes, highlighting the trade-offs between phage and antibiotic resistance^47^. These examples together highlight the rapid turnover of the resistome in *Kpn*, and could help explain the prevalence and ecological success of the species. Beyond the genomic trade-off by which plasmid-borne prophages are streamlined under strong selection pressures (growth/phages), our data also demonstrates that such evolutionary can channel populations toward trade-offs that may be strategically used to restore antibiotic susceptibility.

Overall, our results show that a costly trait can be maintained when its benefits extend beyond the producing cell to other members of the lineage, and help explain the plasticity and high turnover of prophages in genomes. More broadly, our study connects kin selection and social evolution with the biology of temperate phages

## MATERIALS AND METHODS

### Strains

The bacterial strain Kpn BJ1 has no inducible or cryptic prophages and was isolated from a liver abscess (ST380) (ENA: SAMEA4968482). Strain ST14 was isolated from the urinary tract following a nosocomial infection (ENA: SAMN22024794). These two strains are two phylogenetically-distant *Klebsiella pneumoniae* and both encode a K2 capsule. They are here referred to as the phage-sensitive (BJ1) and the polylysogen (ST14). ST14 produces multiple infectious virions for which the BJ1 is known to be sensitive (Supplementary file 1; ^15^). Bacteria were grown at 37°C in Luria-Bertani (LB) agar plates or in 4 mL of liquid broth under vigorous shaking (250 rpm). To avoid terminology confusions, we use the lineage-level benefit when referring to advantages to ST14 cells, rather than population-level or group-level benefits.

### Coevolution experiment

The experimental evolution set-up was previously described in detail ^15^. Briefly, three clones from each strain were used to inoculate overnight cultures, which were then diluted at 1:100 and used to initiate the three independent mixed populations in a ratio 1:1, in a final volume of 4 mL. Each of the three mixed populations evolved in three different environments: (i) LB, (ii) LB supplemented with 0.2% citrate, and (iii) LB with mitomycin C (MMC, 0.1 μg/mL). Cultures were allowed to grow for 24 hours at 37°C and diluted again to 1:100 in fresh media. This was repeated for 30 days. Glycerol stocks of the entire population were frozen every 5 days. Chloramphenicol (30 μg/ml) and trimethoprim (100 μg/ml) were used to select for strain BJ1 and ST14, respectively throughout the evolution experiment.

### Growth curves

Each independent clone was allowed to grow overnight in LB. 200 μL of diluted (1:100) overnight cultures in their evolution environment (either plain LB or supplemented with citrate or with MMC) were distributed in 96-well plates (three wells per clone). Growth was then monitored for 16 hours on a TECAN Spark microtiter plate reader.

### Capsule quantification

The bacterial capsule was extracted as described in Domenico et al., 1989^48^ using 1% Zwittergent 3–14 detergent in 100 mM citric acid (pH 2.0) and heated at 56°C. The extracted capsules were then washed twice in 100% and 70% of absolute ethanol. For each clone, 500 μL of an overnight culture adjusted to OD_600nm_= 2 was measured. Polysaccharides were then quantified by measuring the amount of uronic acid, as described in Blumenkrantz and Asboe-Hansen,1973^49^. The uronic acid concentration in each sample was determined from a standard curve of glucuronic acid.

### PEG-precipitation of phages

Individual clones were grown for in their evolutionary treatment for 24 hours and then centrifuged at 4000 rpm. The supernatant was filtered through 0.22 μm and mixed with chilled PEG-NaCl 5X (PEG 8000 20% and 2.5M of NaCl). Phages were allowed to precipitate for at least 15 min and pelleted by centrifugation 10 min at 13,000 rpm at 4 °C. The pellets were dissolved in TBS (Tris Buffer Saline, 50 mM Tris-HCl, pH 7.5, 150 mM NaCl). Phage concentrations were then determined by plaque forming units on the ancestral prey, strain BJ1.

### Plaque forming units (PFU)

To produce a lysate, each independent clone was grown overnight in its evolutionary environment. Cultures were then centrifuged at 4000 rpm and the supernatant was filtered through 0.22 μm. The presence of competitor did not significantly alter phage production^15^. To quantify the amount of phage produced by each clone, the strain BJ1 (prey) was grown overnight and then diluted 1:100 and allowed to grow until OD_600nm_= 0.8. Two hundred and fifty μL of the culture was mixed with 3 mL of top agar (0.7% agar) and poured into LB plates prewarmed at 37°. Once the plates were dry, lysates were serially diluted in PBS and spotted. Plates were left overnight at room temperature or at 37° for 4 hours. Phage plaques were then counted.

### Microscopy

End-point clones of the sequenced ST14 strain, evolved for 30 days under different environmental conditions, were imaged using phase-contrast Overnight cultures were grown in LB medium at 37°C under shaking conditions (250 rpm). Microscope slides were cleaned with ethanol. 0.6 µL of each overnight culture was deposited by capillarity on 1% agar pads, diluted in M9 medium. Images were captured using an inverted microscope at 100× magnification to visualize cell morphology. Scale bars and image resizing was performed using Fiji v1.0^50^ for macOS.

### Whole genome sequencing

The genomes of end-point clones as well as the ancestral clone were extracted using the Quiagen Genome Extraction Kit prior to Illumina sequencing*. i) Variant calling*. The genome of the ancestral ST14 strain was resequenced using Oxford Nanopore technology and assembled using *Flye* 2.9.6-b1802^51^. It was then corrected using *polypolish* 0.6.0^52^ using Illumina short-read data^53^. To determine genetic changes in representative evolved clones, these were sequenced and compared with ancestral genotype using *breseq* v.0.35.7 ^54^ and *snippy* 4.6.0 (https://github.com/tseemann/snippy) with default parameters. ii) *Phage coverage.* We mapped the reads obtained from Illumina to phage genomes identified by Phastest, using *bwa* (Burrows–Wheeler Aligner) v 0.7.17^55^ with the default parameters. To extract the coverage, we converted the output SAM files to a sorted BAM file using SAMtools ^56^

### Statistic modeling

All statistics were performed in R with tidyverse (v4.3.2). Graphs were generated with ggplot2 package. **Evolutionary trade-offs**. To test whether the cost of increased phage production was reflected in bacterial yield, we fitted a generalized linear model (GLM) using the glm function from the stats package. Model significance was assessed from the fitted regression coefficient. **Competitive fitness model.** To evaluate the relationship between phage production and relative competitive fitness (*W*), we fitted both a quadratic regression model and a generalized additive model (GAM). The quadratic model included linear and squared terms for phage production. The models included the nine evolved populations and three ancestral observations to define the broader fitness landscape; the same qualitative relationship was observed when considering evolved populations alone. Although the quadratic model supported curvature, 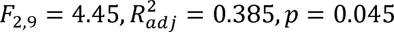, model comparison using Akaike’s Information Criterion (AIC) favored the GAM. GAM was calculated using the mgcv package (v1.9), with a basis dimension capped at *k* = 3 allowing for 2 degrees of freedom, to restrict model over-smoothing and enforce conservative curve complexity. Model adequacy was evaluated using adjusted *R*^2^ 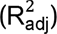, total deviance explained, and approximate -tests for the smooth term. **Bootstrapping evolutionary optimum**. To quantify uncertainty in the evolutionary optimum, we performed 2,000 bootstrap resamples of the observations. The 95% bootstrap confidence interval for the optimum was defined by the 2.5th and 97.5th percentiles of the resulting distribution of peak values.

## ACKNOWLEDGEMENTS

We thank Benoit Pons and Julie Le Bris for critical reading of the manuscript, as well as Arnaud Gutierrez for scientific discussions. We also thank Jerome Rech and Sara Díaz Díaz for their help using the microscope and treating the images. We are grateful to Eduardo P.C. Rocha for his guidance and support during the evolution experiment carried out in his laboratory.

## AUTHOR CONTRIBUTIONS

Olaya Rendueles, Conceptualization, Resources, Data curation, Formal analysis, Supervision, Funding acquisition, Investigation, Visualization, Methodology, Writing - original draft, Project administration, Writing – review and editing.

Teodora Simona Lungu and Louise Badruna-Experiments and data analysis

## SUPPLEMENTARY FIGURE LEGENDS

**Figure S1. Growth of individual clones across time**. Growth is tested both by the area under the curve (AUC) and the maximum yield of a population. Results are presented as relative to their ancestor (dotted line). Each grey dot represents the mean of an independent clone isolated from an independently evolving population. Three different clones from each population at different times points as tested at least in triplicate. The colored dots represent the average of all clones from all populations. Kruskal-Wallis (K-W) tests to compare the effect of evolutionary treatment are shown at the top. Results of one-sample t-tests for differences against the ancestor are shown at the bottom (ns p>0.05, * p <0.05, ** p <0.01 and *** p < 0.001).

**Figure S2. Growth parameters of clones isolated from independently evolving populations. A.** Generation time of each clone relative to the ancestor (μ = 1) was measured in their respective evolutionary environments during the one-hour period of maximum growth. Three different isolated clones from each population at day 30 were tested. Each grey point indicates the average of three independent biological replicates (ns p>0.05, * p <0.05, ** p <0.01 and *** p < 0.001). **B.** Maximum optical density as measured by absorbance at 600nm of MMC-evolved clones and grown in MMC, compared to the ancestor in the absence of induction (dashed line). Clones above the line represent those which outperform the ancestor despite MMC-induction (N=8, 5 of which, significantly so). **C.** Area under the curve using the *trapz* formula from the pracma package of all clones evolved in MMC compared to the ancestor (dashed line) in the absence of induction.

**Figure S3. Phage production of evolved clones.** Results are presented as relative to their ancestor (dotted line). Each grey dot represents the mean of an independent clone isolated from each independently evolving population. The y axis represents the log_10_ of the fold change. Results of one-sample t-tests for differences against the ancestor are shown at the bottom (ns p>0.05, * p <0.05, ** p <0.01 and *** p < 0.001).

**Figure S4. Relative changes in growth versus phage production in evolved clones. A. Trajectories of evolutionary change over time across environments.** Each point represents the average phenotype of three clones (one from each independent population) per evolutionary treatment. Relative changes in growth and phage production were calculated as differences from the ancestral reference (set to 1), based on area under the growth curve (Figure 1) and PFU/mL (Figure 2B), respectively. Dotted lines trace the evolutionary trajectories over time (days 0, 10, 20, 30), with point size increasing with time. Note: PFU/mL changes are represented on a log scale. Error bars are not shown for visibility purposes. **B.** Same as A, but each population and each time point plotted independently. Populations adapted in a step-wise manner with large changes within the first ten days in phage production (large vertical changes), and then optimization of growth (exploration of horizontal space).

**Figure S5. Infection of unpurified lysates from evolved clones. A.** Phages evolved in citrate were tested for their ability to infect even in the presence of citrate. PEG-purified lysates from a representative colony from the end-point population (day 30) were used to infect a population BJ1 at an MO1 of 0,01 in the presence (dark dots) and absence (open dots) of citrate. Both the area under the curve (AUC) and the maximum absorbance (ODmax) reached by the population is shown. Lysate from the ancestor (ST14) is shown as a control. Large dots indicate mean and standard deviation, whereas individual biological replicates are shown in grey. Statistics correspond to a pairwise t-test (* p-value <0.05, ** p-value <0.01 and *** p-value < 0.001). **B.** Lysates from representative individual clones evolved in citrate and isolated at different time points were filtered and added to growing cultures of ancestral prey in the presence or absence of 0.2% citrate (dashed line). Prey was also grown without phage as a control. **C.** The ability of end-point lysates to infect several resistant clones which emerged during the first days of the experiments was assessed in evolutionary conditions. Unpurified ancestral and evolved lysates (ST14 and evolved, respectively) were tested alongside the control condition (plain media). Each resistant clone was tested against a lysate from a representative evolved ST14 clone from the same population. Each panel corresponds to a different resistant clone, or their ancestor (Kpn BJ1). Large dots indicate mean and standard deviation, whereas individual biological replicates are shown in grey. Statistics correspond to pairwise t-tests (ns p-value >0.05, * p-value <0.05, ** p-value <0.01 and *** p-value < 0.001).

**Figure S6. Cell segmentation defects of clones isolated at day 10.** Close-up images of representative micrographs from clones isolated at day 10, from each population from different environmental conditions. Overnight cultures were grown in LB and 5µl of culture were deposited on a microscope slide and pictures were taken with an inverted objective with magnification of x100. Cell segmentation defaults can be observed in some, but not all populations. Scale bar,5 µm.

**Figure S7. Inversion on phage 4 of ST14.** The inverted sequence is highlighted in bright yellow, and is located after a DNA invertase. The annotation was generated using Phastest 3.0.

**Figure S8. Number of reads mapping each phage divided by the total length of the phage in ancestral cultures not induced (A) and induced with MMC (B).**

## SUPPLEMENTARY TABLE LEGENDS

**Table S1. Statistics for growth parameters**. P-values correspond to Wilcoxon tests differences from 1 (ancestral value). Italics highlight those that are statistically significant. The average value of three independent biological replicates were taken into account for each clone.

**Table S2. Mutations in end-point evolved clones compared to their ancestor.** Mutations were mapped against the complete genome of the ancestor using *breseq*, with default settings. Short nucleotide polymorphisms were confirmed independently using *snippy.* Mutations in TatC were confirmed by PCR and Sanger sequencing. ‘na_prot’ indicates a non-annotated protein. Mutations in blue indicate non-synonymous mutations, red indicates premature stop codon and green indicate synonymous mutations. Of note, intergenic mutation at position 4,229,344 appears in all populations. Mutation in phage regions are presented in Table S4.

**Table S3. Phages identified in ancestral strain ST14**. Phages were determined using PHASTEST 3.0. The degree of completeness determined by PHASTEST is prediction of whether the region contains an intact or incomplete prophage based on several criteria including phage protein that match the phage protein database, presence of attachment sites, the number of tRNAs, rRNAs, tmRNA and ORFs in the region. Similarity shows the most common phage in the database, as determined by the phage with the highest number of proteins most similar to those in the region.

**Table S4. Mutations identified in phage regions.** In all evolved clones, a single nucleotide insertion in position 2,625,394, is observed. This corresponds to a tail fiber protein of phage #3. Another single nucleotide insertion in phage #4 is observed in the inverted region also in a fiber protein.

## REFERENCES

1. Hibbing, M. E., Fuqua, C., Parsek, M. R. & Peterson, S. B. Bacterial competition: surviving and thriving in the microbial jungle. Nat. Rev. Microbiol. 8, 15–25 (2010).

2. Rendueles, O. & Ghigo, J.-M. Multi-species biofilms: how to avoid unfriendly neighbors. FEMS Microbiol. Rev. 36, 972–989 (2012).

3. Hawlena, H., Bashey, F. & Lively, C. M. Bacteriocin-mediated interactions within and between coexisting species. Ecol. Evol. 2, 2521–2526 (2012).

4. Peterson, S. B., Bertolli, S. K. & Mougous, J. D. The Central Role of Interbacterial Antagonism in Bacterial Life. Curr. Biol. CB 30, R1203–R1214 (2020).

5. Granato, E. T., Meiller-Legrand, T. A. & Foster, K. R. The Evolution and Ecology of Bacterial Warfare. Curr. Biol. CB 29, R521–R537 (2019).

6. Freilich, S. et al. Competitive and cooperative metabolic interactions in bacterial communities. Nat. Commun. 2, 589 (2011).

7. Schlechter, R. O., Kear, E. J., Bernach, M., Remus, D. M. & Remus-Emsermann, M. N. P. Metabolic resource overlap impacts competition among phyllosphere bacteria. ISME J. 17, 1445–1454 (2023).

8. Foster, K. R. & Bell, T. Competition, not cooperation, dominates interactions among culturable microbial species. Curr. Biol. CB 22, 1845–1850 (2012).

9. Touchon, M., Bernheim, A. & Rocha, E. P. Genetic and life-history traits associated with the distribution of prophages in bacteria. ISME J. 10, 2744–2754 (2016).

10. West, S. A., Griffin, A. S. & Gardner, A. Evolutionary Explanations for Cooperation. Curr. Biol. 17, R661–R672 (2007).

11. Hamilton, W. D. The genetical evolution of social behaviour. I. J. Theor. Biol. 7, 1– 16 (1964).

12. Burns, N., James, C. E. & Harrison, E. Polylysogeny magnifies competitiveness of a bacterial pathogen in vivo. Evol. Appl. 8, 346–351 (2015).

13. Niehus, R., Oliveira, N. M., Li, A., Fletcher, A. G. & Foster, K. R. The evolution of strategy in bacterial warfare via the regulation of bacteriocins and antibiotics. eLife 10, e69756 (2021).

14. de Sousa, J. A. M., Buffet, A., Haudiquet, M., Rocha, E. P. C. & Rendueles, O. Modular prophage interactions driven by capsule serotype select for capsule loss under phage predation. ISME J. 14, 2980–2996 (2020).

15. Rendueles, O., de Sousa, J. A. & Rocha, E. P. Competition between lysogenic and sensitive bacteria is determined by the fitness costs of the different emerging phage-resistance strategies. eLife 12, e83479 (2023).

16. Shafia, F. & Thompson, T. L. Calcium ion requirement for proliferation of bacteriophage phi mu-4. J. Bacteriol. 88, 293–296 (1964).

17. Otsuji, N., Sekiguchi, M., Iijima, T. & Takagi, Y. Induction of phage formation in the lysogenic Escherichia coli K-12 by mitomycin C. Nature 184(Suppl 14), 1079–1080 (1959).

18. Frazão, N. et al. Two modes of evolution shape bacterial strain diversity in the mammalian gut for thousands of generations. Nat. Commun. 13, 5604 (2022).

19. Bobay, L.-M., Touchon, M. & Rocha, E. P. C. Pervasive domestication of defective prophages by bacteria. Proc. Natl. Acad. Sci. U. S. A. 111, 12127–12132 (2014).

20. Bailey, Z. M., Igler, C. & Wendling, C. C. Prophage maintenance is determined by environment-dependent selective sweeps rather than mutational availability. Curr. Biol. CB 34, 1739–1749.e7 (2024).

21. Khan, A., Burmeister, A. R. & Wahl, L. M. Evolution along the parasitism-mutualism continuum determines the genetic repertoire of prophages. PLoS Comput. Biol. 16, e1008482 (2020).

22. Levin, P. A. & Janakiraman, A. Localization, Assembly, and Activation of the Escherichia coli Cell Division Machinery. EcoSal Plus 9, eESP00222021 (2021).

23. Cameron, T. A. & Margolin, W. Insights into the assembly and regulation of the bacterial divisome. Nat. Rev. Microbiol. 22, 33–45 (2024).

24. Mueller, E. A. et al. The active repertoire of *Escherichia coli* peptidoglycan amidases varies with physiochemical environment. Mol. Microbiol. 116, 311–328 (2021).

25. Wilksch, J. J. et al. MrkH, a Novel c-di-GMP-Dependent Transcriptional Activator, Controls Klebsiella pneumoniae Biofilm Formation by Regulating Type 3 Fimbriae Expression. PLOS Pathog. 7, e1002204 (2011).

26. Belfort, M. & Wulff, D. The Roles of the Lambda *c* III Gene and the *Escherichia coli* Catabolite Gene Activation System in the Establishment of Lysogeny by Bacteriophage Lambda. Proc. Natl. Acad. Sci. 71, 779–782 (1974).

27. Noble, J. A. et al. The Escherichia coli hflA locus encodes a putative GTP-binding protein and two membrane proteins, one of which contains a protease-like domain. Proc. Natl. Acad. Sci. 90, 10866–10870 (1993).

28. Grant, J. R. et al. Proksee: in-depth characterization and visualization of bacterial genomes. Nucleic Acids Res. 51, W484–W492 (2023).

29. Wishart, D. S. et al. PHASTEST: faster than PHASTER, better than PHAST. Nucleic Acids Res. 51, W443–W450 (2023).

30. Alcock, B. P. et al. CARD 2023: expanded curation, support for machine learning, and resistome prediction at the Comprehensive Antibiotic Resistance Database. Nucleic Acids Res. 51, D690–D699 (2023).

31. Touchon, M., Moura De Sousa, J. A. & Rocha, E. P. Embracing the enemy: the diversification of microbial gene repertoires by phage-mediated horizontal gene transfer. Curr. Opin. Microbiol. 38, 66–73 (2017).

32. Nanda, A. M., Thormann, K. & Frunzke, J. Impact of Spontaneous Prophage Induction on the Fitness of Bacterial Populations and Host-Microbe Interactions. J. Bacteriol. 197, 410–419 (2015).

33. Bossi, L., Fuentes, J. A., Mora, G. & Figueroa-Bossi, N. Prophage contribution to bacterial population dynamics. J. Bacteriol. 185, 6467–6471 (2003).

34. Vriezen, J. A. C., Valliere, M. & Riley, M. A. The Evolution of Reduced Microbial Killing. Genome Biol. Evol. 1, 400–408 (2009).

35. Banuett, F. & Herskowitz, I. Identification of polypeptides encoded by an Escherichia coli locus (hflA) that governs the lysis-lysogeny decision of bacteriophage lambda. J. Bacteriol. 169, 4076–4085 (1987).

36. Abedon, S. T. Lysis from without. Bacteriophage 1, 46–49 (2011).

37. Buffet, A., Rocha, E. P. C. & Rendueles, O. Nutrient conditions are primary drivers of bacterial capsule maintenance in Klebsiella. Proc. Biol. Sci. 288, 20202876 (2021).

38. Beamud, B. et al. Genetic determinants of host tropism in Klebsiella phages. Cell Rep. 42, 112048 (2023).

39. Le Bris, J., Varet, H., Rocha E.P.C., and Olaya Rendueles. Plug-and-play dynamics of capsule serotype swapping in Klebsiella spp. BioRXiv (2025).

40. Scanlan, P. D. et al. Coevolution with Bacteriophages Drives Genome-Wide Host Evolution and Constrains the Acquisition of Abiotic-Beneficial Mutations. Mol. Biol. Evol. 32, 1425–1435 (2015).

41. Gittrich, M. R. et al. Cross-family and phage-specific gene requirements for Klebsiella infection revealed by scalable RB-TnSeq genetic screens. PLOS Genet. 22, e1012233 (2026).

42. Ouyang, R., Ongenae, V., Muok, A., Claessen, D. & Briegel, A. Phage fibers and spikes: a nanoscale Swiss army knife for host infection. Curr. Opin. Microbiol. 77, 102429 (2024).

43. Sanjuán, R. The Social Life of Viruses. Annu. Rev. Virol. 8, 183–199 (2021).

44. Dunai, A. et al. Rapid decline of bacterial drug-resistance in an antibiotic-free environment through phenotypic reversion. eLife 8, e47088 (2019).

45. Lamrabet, O., Martin, M., Lenski, R. E. & Schneider, D. Changes in Intrinsic Antibiotic Susceptibility during a Long-Term Evolution Experiment with Escherichia coli. mBio 10, e00189–19 (2019).

46. Matthews, A. C., Lehtinen, S. & Dimitriu, T. Plasmid streamlining drives the extinction of antibiotic resistance plasmids under selection for horizontal transmission. PLOS Biol. 23, e3003564 (2025).

47. Majkowska-Skrobek, G., et al. The evolutionary trade-offs in phage-resistant *KLEBSIELLA PNEUMONIAE* entail cross-phage sensitization and loss of multidrug resistance. Environ. Microbiol. 23, 7723–7740 (2021).

48. Domenico, P., Salo, R. J., Cross, A. S. & Cunha, B. A. Polysaccharide capsule-mediated resistance to opsonophagocytosis in Klebsiella pneumoniae. Infect. Immun. 62, 4495–4499 (1994).

49. Blumenkrantz, N. & Asboe-Hansen, G. New method for quantitative determination of uronic acids. Anal. Biochem. 54, 484–489 (1973).

50. Schindelin, J. et al. Fiji: an open-source platform for biological-image analysis. Nat. Methods 9, 676–682 (2012).

51. Kolmogorov, M., Yuan, J., Lin, Y. & Pevzner, P. A. Assembly of long, error-prone reads using repeat graphs. Nat. Biotechnol. 37, 540–546 (2019).

52. Wick, R. R. & Holt, K. E. Polypolish: Short-read polishing of long-read bacterial genome assemblies. PLOS Comput. Biol. 18, e1009802 (2022).

53. Blin, C., Passet, V., Touchon, M., Rocha, E. P. C. & Brisse, S. Metabolic diversity of the emerging pathogenic lineages of Klebsiella pneumoniae. Environ. Microbiol. 19, 1881–1898 (2017).

54. Deatherage, D. E. & Barrick, J. E. Identification of Mutations in Laboratory-Evolved Microbes from Next-Generation Sequencing Data Using breseq. in Engineering and Analyzing Multicellular Systems: Methods and Protocols (eds Sun, L. & Shou, W.) 165–188 (Springer, New York, NY, 2014). doi:10.1007/978-1-4939-0554-6_12.

55. Langmead, B. & Salzberg, S. L. Fast gapped-read alignment with Bowtie 2. Nat. Methods 9, 357–359 (2012).

56. Li, H. et al. The Sequence Alignment/Map format and SAMtools. Bioinformatics 25, 2078–2079 (2009).

