## Supplementary Figures for "Environment-dependent evolution drives divergent adaptive strategies balancing parasite costs and competitive benefits"

### SUPPLEMENTARY FIGURES for Environment-dependent evolution drives divergent adaptive strategies and parasite dynamics in a minimal community

Teodora Simona Lungu[1], Louise Badruna[1] and Olaya Rendueles[1,2#]

[1]Laboratoire de Microbiologie et Génétique Moléculaires (LMGM), CNRS UMR5100, Centre de Biologie Intégrative (CBI), Université de Toulouse, CNRS, Université Toulouse III - Paul Sabatier (UT3), Toulouse, France

[2]Institut Pasteur, Université Paris Cité, CNRS UMR3525, Microbial Evolutionary Genomics, Paris 75015, France.

ORCID: OR, 0000-0002-6648-1594

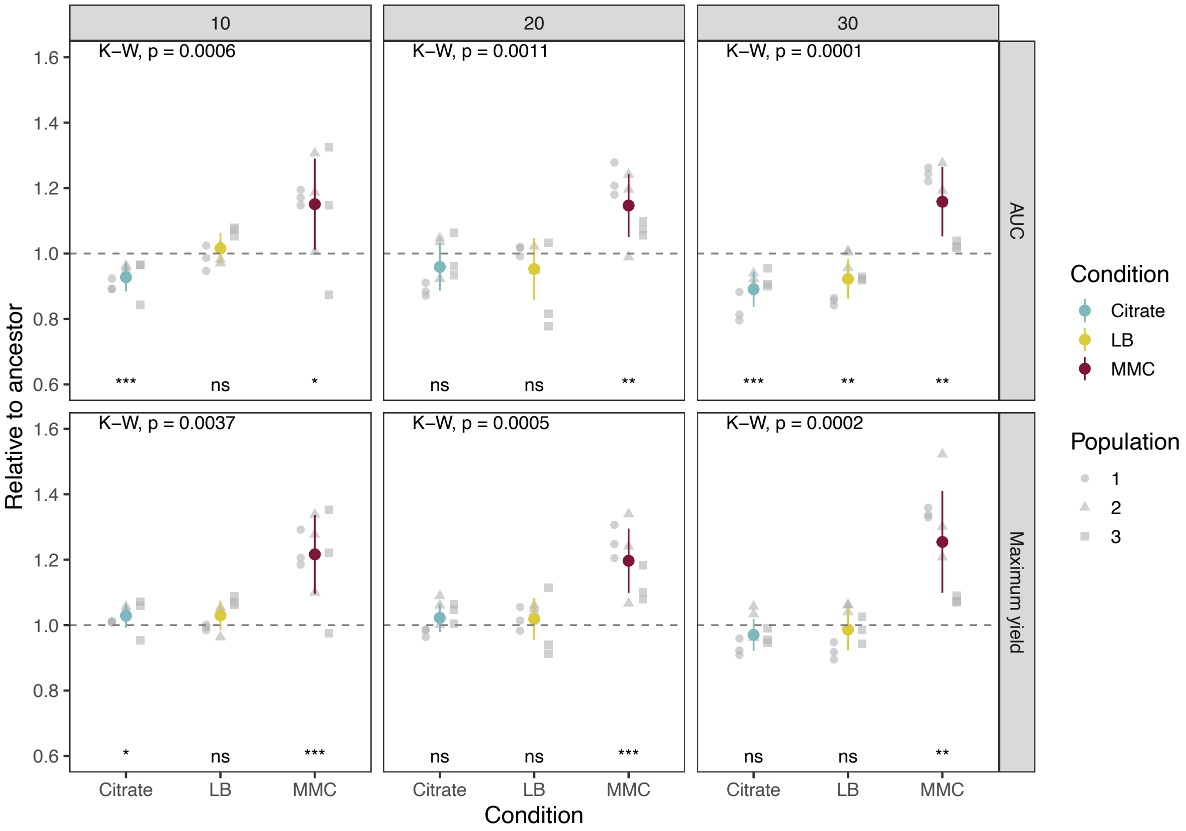


**Figure S1. Growth of individual clones across time**. Growth is tested both by the area under the curve (AUC) and the maximum yield of a population. Results are presented as relative to their ancestor (dotted line). Each grey dot represents the mean of an independent clone isolated from an independently evolving population. Three different clones from each population at different times points as tested at least in triplicate.

The colored dots represent the average of all clones from all populations. Kruskal-Wallis (K-W) tests to compare the effect of evolutionary treatment are shown at the top. Results of one-sample t-tests for differences against the ancestor are shown at the bottom (ns p>0.05 ,* p <0.05, ** p <0.01 and *** p < 0.001).

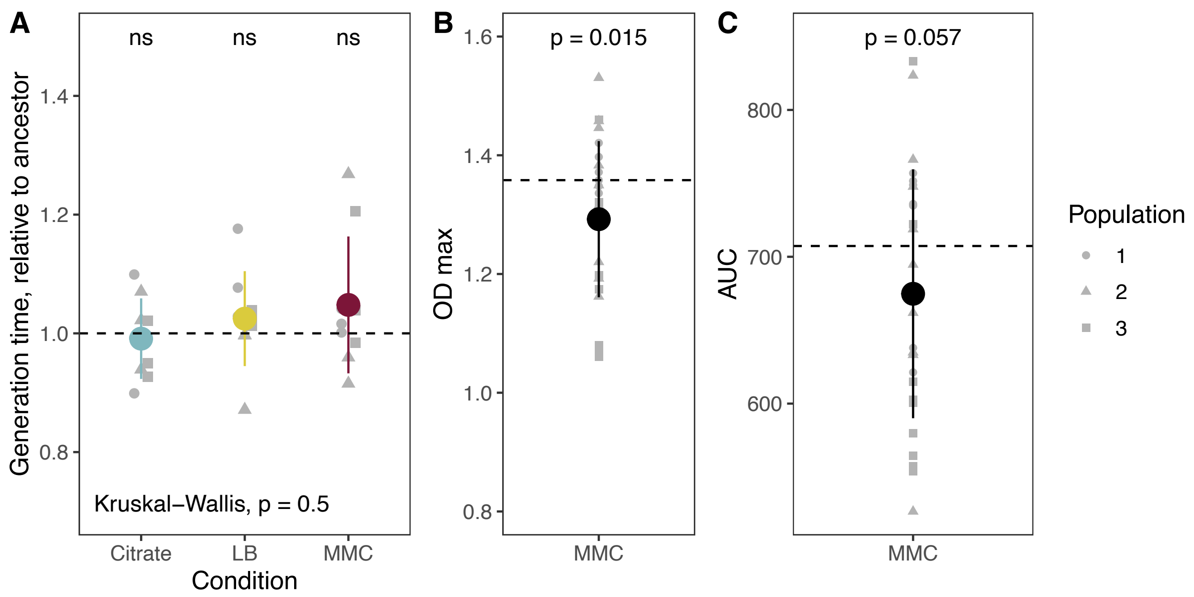


**Figure S2. Growth parameters of clones isolated from independently evolving populations. A.** Generation time of each clone relative to the ancestor (μ = 1) was measured in their respective evolutionary environments during the one-hour period of maximum growth. Three different isolated clones from each population at day 30 were tested. Each grey point indicates the average of three independent biological replicates (ns p>0.05 ,* p <0.05, ** p <0.01 and *** p < 0.001). **B.** Maximum optical density as measured by absorbance at 600nm of MMC-evolved clones and grown in MMC, compared to the ancestor in the absence of induction (dashed line). Clones above the line represent those which outperform the ancestor despite MMC-induction (N=8, 5 of which, significantly so). **C.** Area under the curve using the *trapz* formula from the pracma package of all clones evolved in MMC compared to the ancestor (dashed line) in the absence of induction.

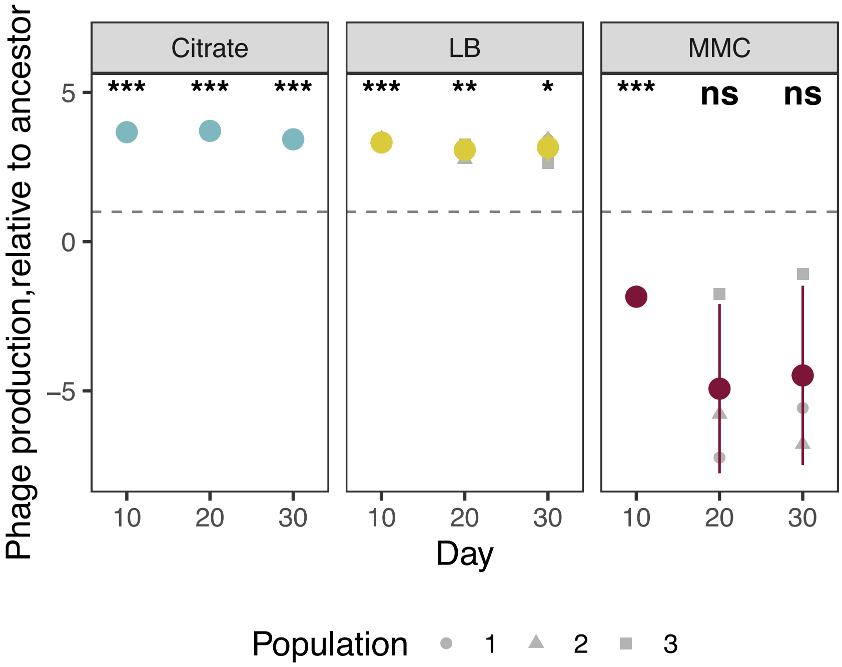


**Figure S3. Phage production of evolved clones.** Results are presented as relative to their ancestor (dotted line). Each grey dot represents the mean of an independent clone isolated from each independently evolving population. The y axis represents the $log_{10}$ of the fold change. Results of one-sample t-tests for differences against the ancestor are shown at the bottom (ns p>0.05 ,* p <0.05, ** p <0.01 and *** p < 0.001).

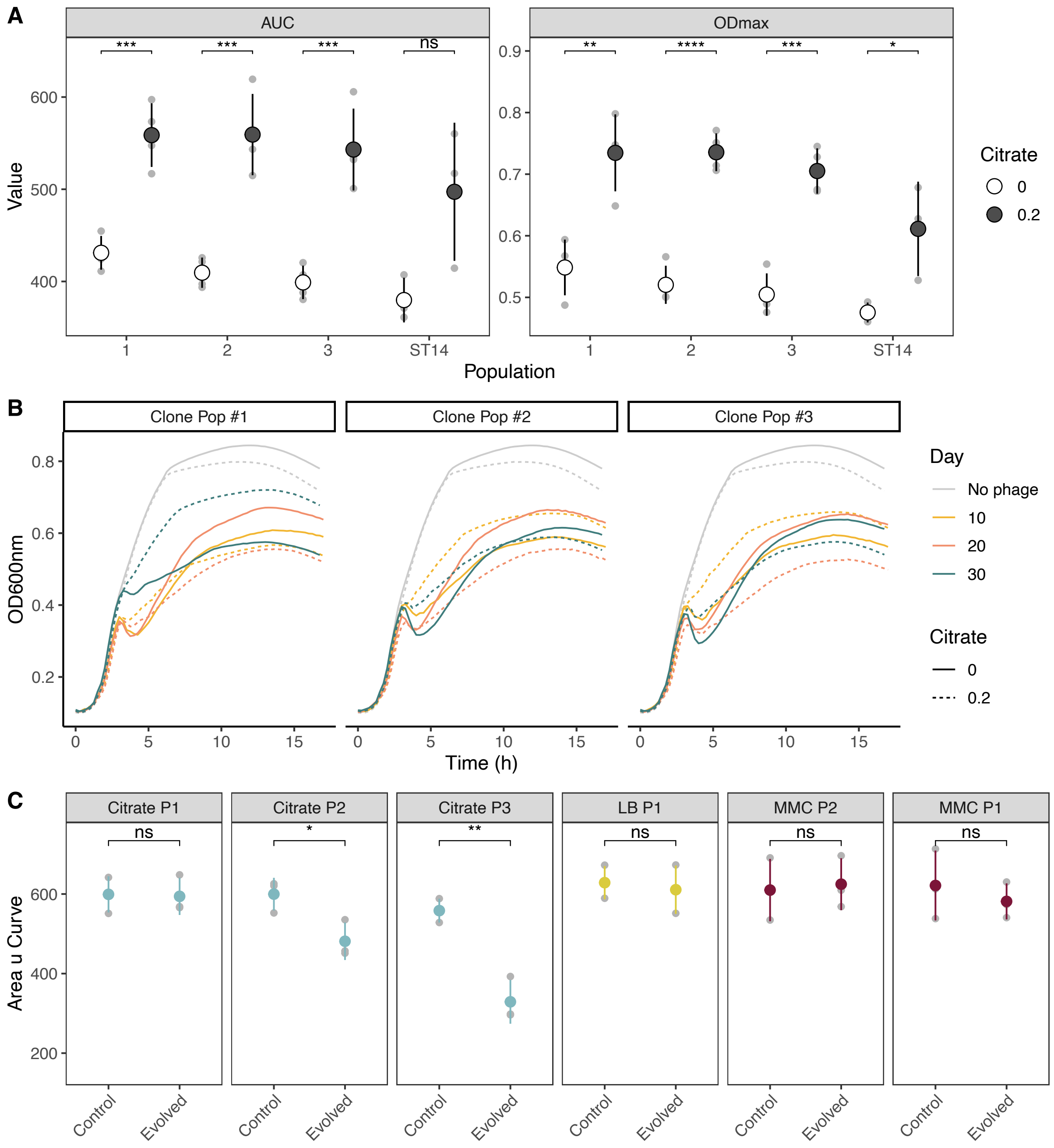


**Figure S4. Infection of unpurified lysates from evolved clones.A.** Phages evolved in citrate were tested for their ability to infect even in the presence of citrate. PEG-purified lysates from a representative colony from the end-point population (day 30) were used to infect a population BJ1 at an MO1 of 0,01 in the presence (dark dots) and absence (open dots) of citrate. Both the area under the curve (AUC) and the maximum absorbance (ODmax) reached by the population is shown. Lysate from the ancestor (ST14) is shown as a control. Large dots indicate mean and standard deviation, whereas individual biological replicates are shown in grey. Statistics correspond to a pairwise t-test (* p-value <0.05, ** p-value <0.01 and *** p-value < 0.001). **B.** Lysates from representative individual clones evolved in citrate and isolated at different time points were filtered and added to growing cultures of ancestral prey in the presence or absence of 0.2% citrate (dashed line). Prey was also grown without phage as a control. **C.** The ability of end-point lysates to infect several resistant clones which emerged during the first days of the experiments was assessed in evolutionary conditions. Unpurified ancestral and evolved lysates (ST14 and evolved, respectively) were tested alongside the control condition (plain media). Each resistant clone was tested against a lysate from a representative evolved ST14 clone from the same population. Each panel corresponds to a different resistant clone, or their ancestor (Kpn BJ1). Large dots indicate mean and standard deviation, whereas individual biological replicates are shown in grey. Statistics correspond to pairwise t-tests (ns p-value >0.05, * p-value <0.05, ** p-value <0.01 and *** p-value < 0.001).

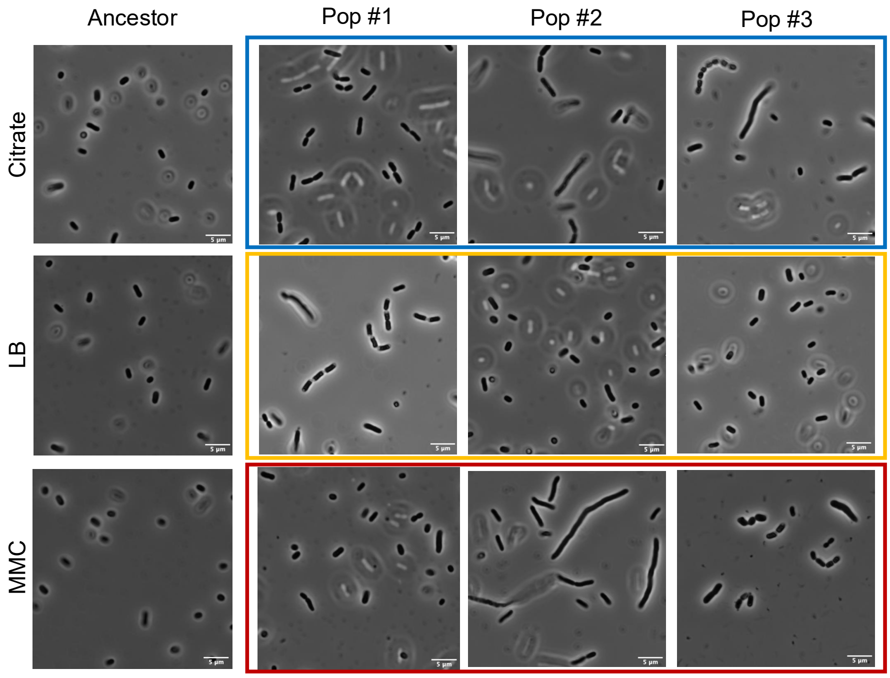


**Figure S5. Cell segmentation defects of clones isolated at day 10.** Close-up images of representative micrographs from clones isolated at day 10, from each population from different environmental conditions. Overnight cultures were grown in LB and 5µl of culture were deposited on a microscope slide and pictures were taken with an inverted objective with magnification of x100. Cell segmentation defaults can be observed in some, but not all populations. Scale bar,5 µm.

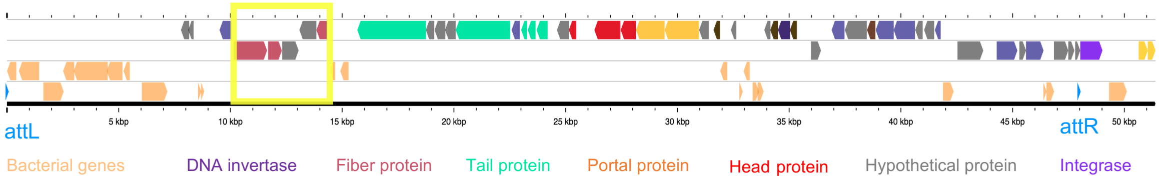


**Figure S6. Inversion on phage 4 of ST14.** The inverted sequence is highlighted in bright yellow, and is located after a DNA invertase. The annotation was generated using Phastest 3.0.

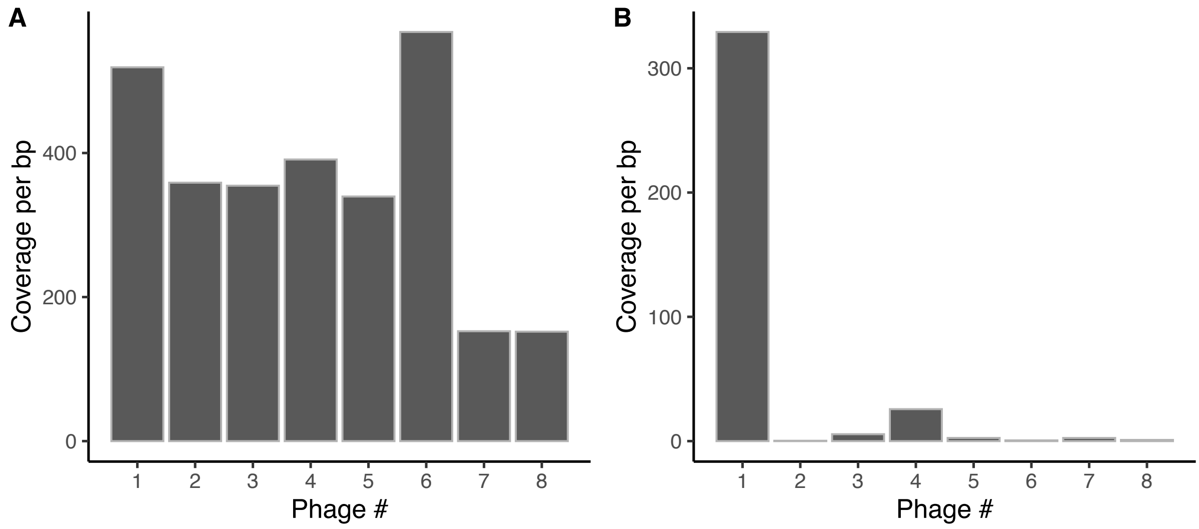


**Figure S7. Number of reads mapping each phage divided by the total length of the phage in ancestral cultures not induced (A) and induced with MMC (B).**

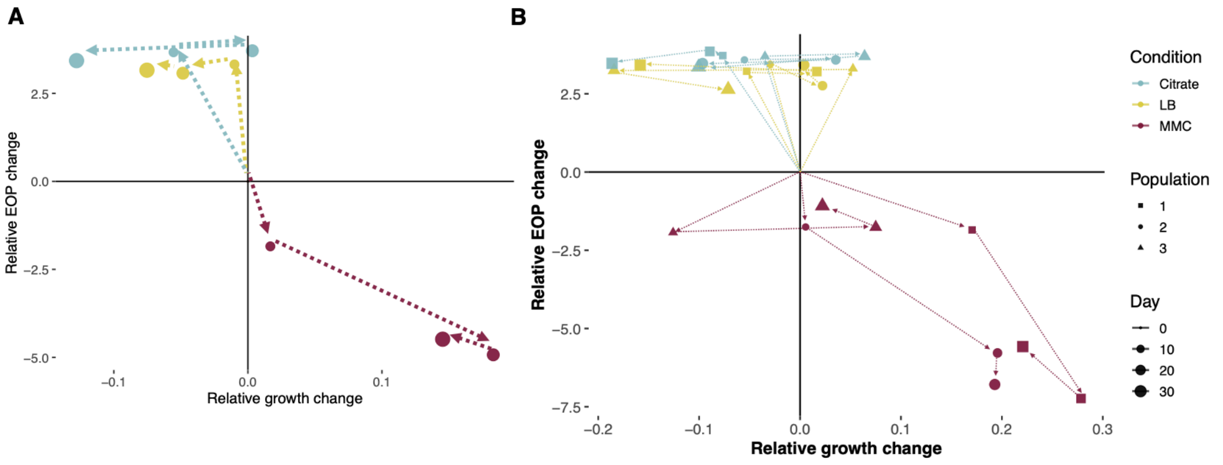


**Figure S8. Relative changes in growth versus phage production in evolved clones. A. Trajectories of evolutionary change over time across environments.** Each point represents the average phenotype of sequenced clones from three independent populations per evolutionary treatment. Relative changes in growth and phage production were calculated as differences from the ancestral reference (set to 1), based on area under the growth curve (Figure 1) and PFU/mL (Figure 2B), respectively. Dotted lines trace the evolutionary trajectories over time (days 0, 10, 20, 30), with point size increasing with time. Note: PFU/mL changes are represented on a $\log_{10}$ scale. Correlation between the change in phage production as measured by PFU/mL and growth as measured by AUC in end-point clones, relative to their common ancestor. **B.** Same as A, but each population and each time point plotted independently.

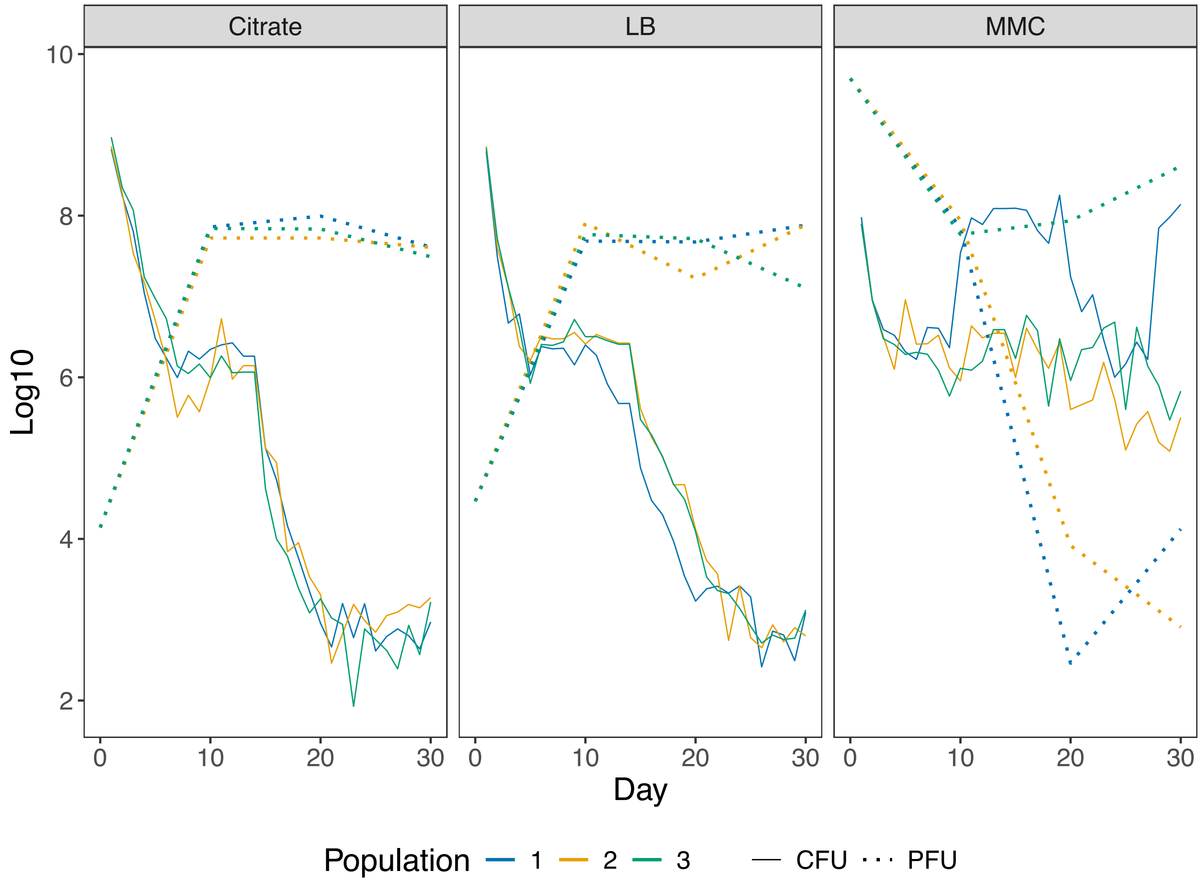


**Figure S9. Comparison of phage abundance and total density of Kpn BJ1 (prey).** Dashed lines correspond to phage populations as measured by PFU/mL on ancestral prey. Solid lines correspond to colony forming units of Kpn BJ1, initially phage-sensitive bacteria in the consortium. Kpn BJ1 was counted every day throughout the evolution experiment. Plaque forming units per mL was assessed at day 10, 20 and 30 after the evolution experiment. Error bars are not shown for visibility purposes.
